# CellChat for systematic analysis of cell-cell communication from single-cell and spatially resolved transcriptomics

**DOI:** 10.1101/2023.11.05.565674

**Authors:** Suoqin Jin, Maksim V. Plikus, Qing Nie

## Abstract

Recent advances in single-cell sequencing technologies offer an opportunity to explore cell-cell communication in tissues systematically and with reduced bias. A key challenge is the integration between known molecular interactions and measurements into a framework to identify and analyze complex cell-cell communication networks. Previously, we developed a computational tool, named CellChat that infers and analyzes cell-cell communication networks from single-cell RNA-sequencing (scRNA-seq) data within an easily interpretable framework. CellChat quantifies the signaling communication probability between two cell groups using a simplified mass action-based model, which incorporates the core interaction between ligands and receptors with multi-subunit structure along with modulation by cofactors. CellChat v2 is an updated version that includes direct incorporation of spatial locations of cells, if available, to infer spatially proximal cell-cell communication, additional comparison functionalities, expanded database of ligand-receptor pairs along with rich annotations, and an Interactive CellChat Explorer. Here we provide a step-by-step protocol for using CellChat v2 that can be used for both scRNA-seq and spatially resolved transcriptomic data, including inference and analysis of cell-cell communication from one dataset and identification of altered signaling across different datasets. The key steps of applying CellChat v2 to spatially resolved transcriptomics are described in detail. The R implementation of CellChat v2 toolkit and tutorials with the graphic outputs are available at https://github.com/jinworks/CellChat. This protocol typically takes around 20 minutes, and no specialized prior bioinformatics training is required to complete the task.

## Introduction

Cell-cell communication orchestrates tissue organization. Recent advances in single-cell genomics offer unprecedented opportunity to systematically explore signaling mechanisms for cell fate decisions and their consequent tissue phenotypes. Using single-cell transcriptomics data and ligand-receptor interaction information from prior knowledge, computational methods have been developed for inferring and analyzing cell-cell communication between groups of cells^1-3^. However, spatial information on cells is inherently lost in single-cell transcriptomics. Given that most cell-cell communication events occur within a short spatial distance, the emerging spatially resolved transcriptomics bring new ways to improve cell-cell communication analysis^4-6^.

### Development of the protocol

Comprehensive and accurate recapitulation of known molecular interactions is crucial for predicting biologically meaningful intercellular communications. We manually curated a literature-supported signaling molecule interaction database called CellChatDB^7^, which takes into account several critical interaction mechanisms that are often neglected. Specifically, CellChatDB not only incorporates multi-subunit structure of ligand-receptor complexes but also accounts for soluble and membrane-bound stimulatory and inhibitory cofactors such as agonists, antagonists and co-receptors (Fig. 1). In addition, CellChat classifies each ligand-receptor pair into one of the functionally related signaling pathways, in order to construct cell-cell communication at a signaling pathway level. In particular, each link of the network is computed by summing the interaction strengths of all associated ligand-receptor pairs. Such information allows the interpretation of inferred intercellular communications at a pathway scale.

**Figure 1:**
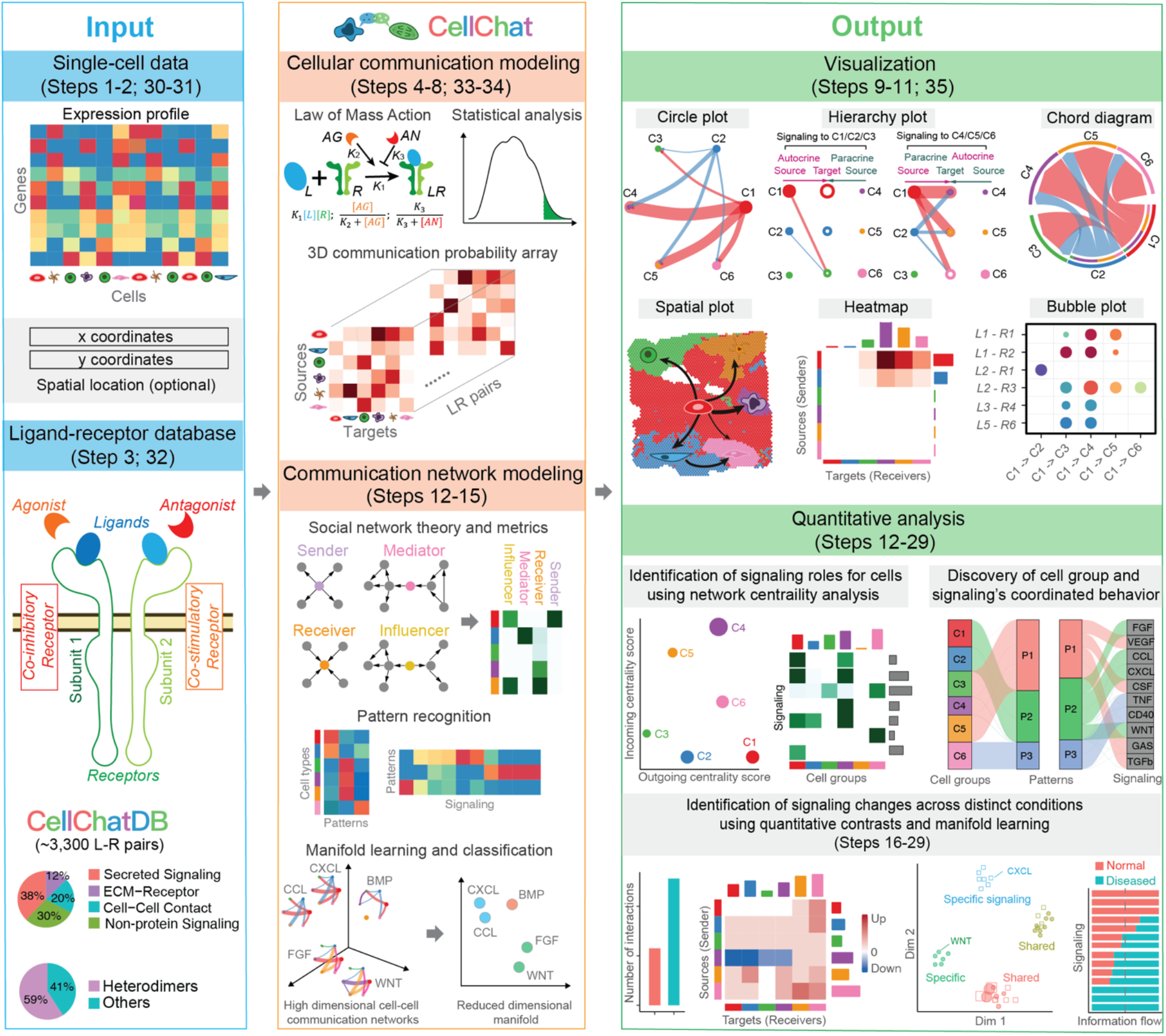
Overview of CellChat along with the Procedure step numbers. **Left:** Required input data and the ligand-receptor interaction database CellChatDB. CellChat’s input data consist of gene expression data and cell group information. When inferring spatially-proximal cell-cell communication, spatial locations of cells are also required. CellChatDB takes into account known composition of the ligand-receptor complexes, including complexes with multimeric ligands and receptors, as well as several cofactor types: soluble agonists, antagonists, co-stimulatory and co-inhibitory membrane-bound receptors. **Middle:** CellChat models the communication probability based on the law of mass action and identifies significant communications using permutation tests. The inferred communication probabilities among all pairs of cell groups across all pairs of ligand-receptor are represented by a three-dimensional array. CellChat analyzes the inferred networks by leveraging social network metrics, pattern recognition methods and manifold learning approaches. **Right:** CellChat offers several intuitive visualization outputs to facilitate data interpretation of different analytical tasks. In addition to analyzing individual dataset, CellChat also delineates signaling changes across different conditions. The Procedure step numbers in different parts are also indicated to link the Procedure sections with the overall scheme.

To quantify communications between two cell groups mediated by a given ligand and its cognate receptor, CellChat leverages the law of mass action to associate each interaction with an interaction score^7^, which is calculated based on the average expression values of a ligand by one cell group and that of a receptor by another cell group, as well as their cofactors (Fig. 1). Here CellChat uses Hill functions in the simplified mass action model to reflect the saturation effect of the ligand-receptor binding. Significant interactions are identified based on a statistical test that randomly permutes the group labels of cells. When inferring cell-cell communication, CellChat computationally scales well with the number of cells and cell groups in the data, as reflected by the observed running time of ∼15 mins on a single cell atlas of adult human skin with ∼300,000 cells (Fig. 2). It should be noted that the number of inferred signaling depends on the method for calculating average gene expression per cell group. The most stringent method “triMean” produces fewer but stronger interactions, while the “truncated mean” method with smaller values of “trim” parameter enables the identification of weak signaling (Fig. 3).

**Figure 2:**
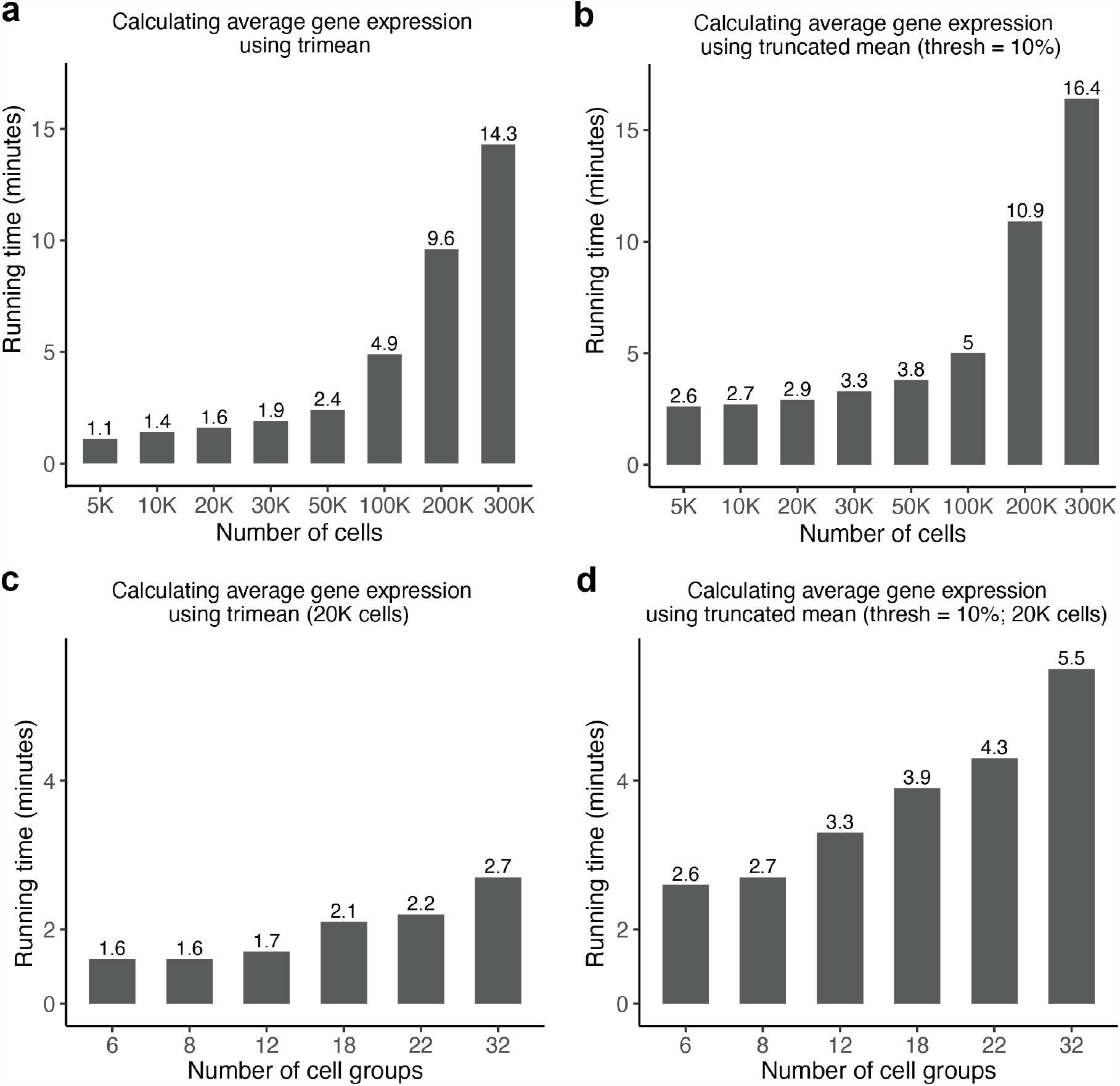
CellChat’s running time with the increase of cell numbers and cell groups. Running time over different cell numbers in the data when calculating average gene expression per cell group using trimean **(a)** or 10% truncated mean **(b)**. Running time over different numbers of cell groups in the data (#20,000 cells) when calculating average gene expression per cell group using trimean **(c)** or 10% truncated mean **(d)**. Here the running time is the total time when running Steps 1-7.

**Figure 3:**
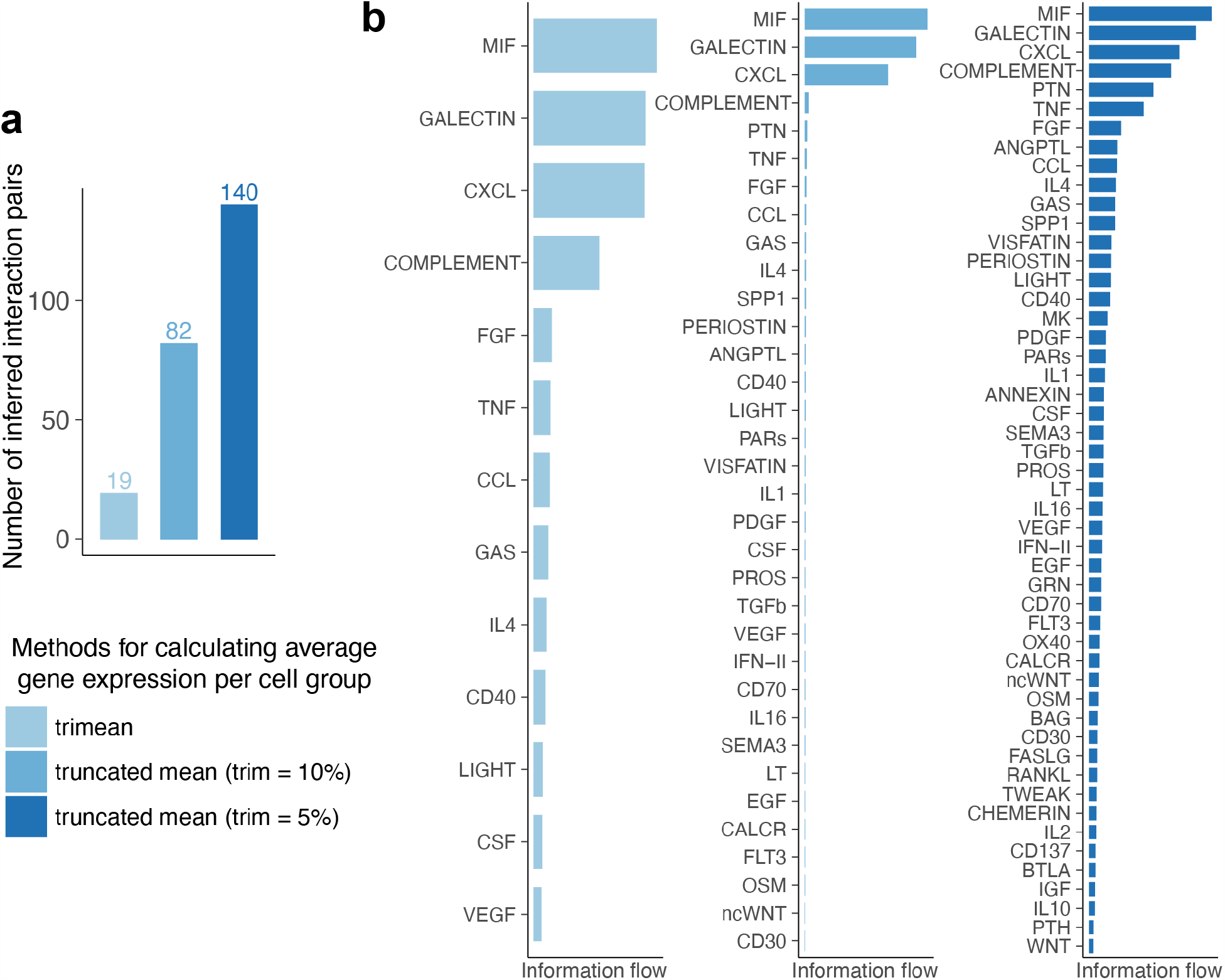
Comparison of the number of inferred ligand-receptor pairs and the identified signaling pathways when using different methods for calculating average gene expression per cell group. **(a)** The number of inferred ligand-receptor pairs when using three different methods for calculating average gene expression per cell group, including trimean, 10% truncated mean and 5% truncated mean. **(b)** The identified signaling pathways when using trimean, 10% truncated mean and 5% truncated mean. The most stringent method “triMean” produces fewer but stronger interactions, while the “truncated mean” method with smaller values of “trim” parameter enables the identification of weak signaling.

To obtain biological insights from a large number of complicated cell-cell communication networks, CellChat employs quantitative analysis and machine learning approaches for the following critical analysis tasks^7^ (Fig. 1). First, to identify critical microenvironment components, CellChat can determine major signaling sources and targets, as well as mediators and influencers within a given signaling network using network centrality analysis. Second, to reveal how cells and signals coordinate together, CellChat can predict key incoming and outgoing signals for specific cell types, as well as coordinated responses among different cell types by leveraging pattern recognition approaches. Outgoing patterns reveal how sender cells (i.e., cells acting as signal source) coordinate with each other, as well as how they coordinate with certain signaling pathways to drive communication. Incoming patterns show how target cells (i.e., cells acting as signal receivers) coordinate with each other to respond to incoming signals. Third, to predict signaling groups with similar communication architecture and interpret the biological functions of poorly studied pathways, CellChat groups signaling pathways by defining similarity measures and performing manifold learning from both functional and topological perspectives^7^.

To identify signaling changes across conditions, CellChat first focuses on the overall signaling changes at the cell population level and then narrows down to altered signaling pathways and ligand-receptor pairs^7-9^ (Fig. 1). At the cell population level, CellChat identifies which interactions between which two cell groups are significantly changed; as well as cell group identities showing significant changes in sending or receiving signaling patterns across conditions. At the level of signaling pathways and ligand-receptor pairs, CellChat identifies altered signaling pathways and ligand-receptor pairs in terms of network architecture and interaction strength by performing joint manifold learning and information flow comparison analysis. To identify statistically significantly upregulated and downregulated ligand-receptor pairs across conditions, CellChat combines cell-cell communication analysis with differential gene expression analysis and quantifies the enrichment of ligand-receptor pairs in a given condition by defining an enrichment score.

Compared to the original CellChat, the updated CellChat v2, enables inference of physically proximal cell-cell communication between interacting cell groups if spatial locations of cells are available (Fig. 4a). Inference of cell-cell communication can be naturally extended to spatial context by first identifying pairs of cells that are physically close to one another to have biologically realistic interactions based on maximal possible molecular interaction/diffusion ranges, and then identifying combinations of cell groups that have enough nearby cell-cell pairs. The diffusive spatial distance of molecules depends on many factors, including molecule size, its covalent modifications, touristy of the spatial tissues, and the molecule’s regulators on the cell membrane and in the extracellular environments. All these factors usually reduce diffusion. For example, large molecules have shorter diffusion distance, leading to more restricted spatial range in diffusion. In CellChat v2, we use the ideal diffusion range in a free medium, that is the maximally allowable transport distance for small diffusive molecules (by default 250 *μm*). In this way, CellChat v2 will not remove any interactions that are spatially plausible. For the contact-dependent signaling, the interaction range is restricted to the nearest neighbors of each cell, such that signaling and target cells are in direct contact.

**Figure 4:**
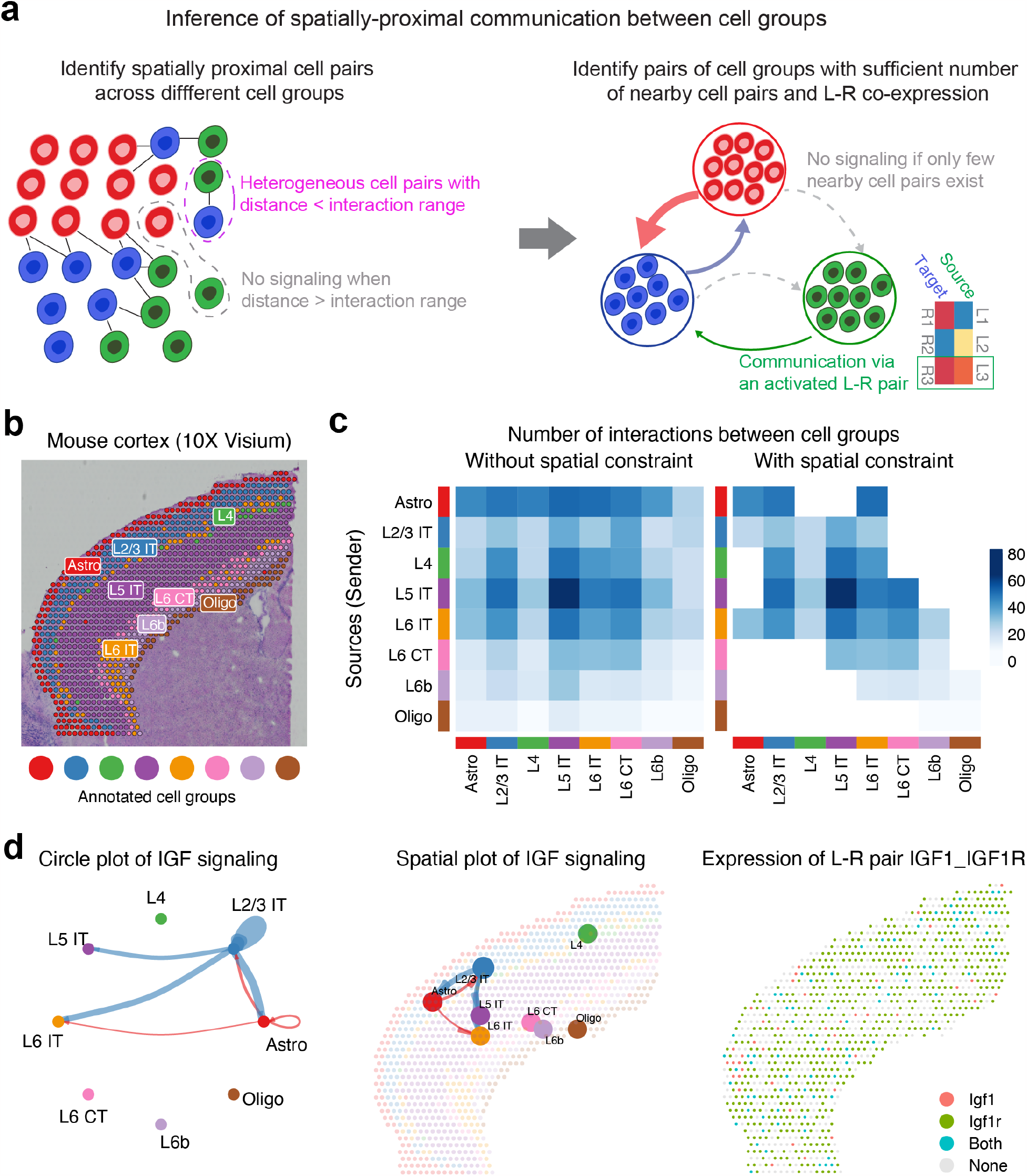
Application of CellChat v2 to spatially resolved transcriptomics. **(a)** CellChat v2 infers spatially-proximal cell-cell communication between adjacent groups of cells, modelled by first identifying pairs of cells that are physically proximal within the maximum assumed interaction/diffusion range, and then identifying pairs of cell groups that have sufficient number of nearby cell pairs. **(b)** An example of spatially resolved transcriptomic data from 10X Visium experiment on mouse cortex. Annotations of each spot are used as cell group information for CellChat analysis. **(c)** Comparison of the number of interactions between pairs of cell groups using spatial constraint against the case without spatial constraint. **(d)** Example visualization outputs for signaling analysis in the spatial context. The typical circle plot shows both autocrine and paracrine signaling interactions, but the spatial plot only shows the paracrine signaling interactions between different cell groups. The feature plot shows the expression distribution of a ligand and its receptor by binarizing their expression in each spot. Each spot is colored based on whether it only expresses the ligand, the receptor or both signaling molecules.

In addition, in CellChat v2 we have: (a) expanded the CellChatDB to include more than 1000 protein and non-protein interactions (e.g., metabolic and synaptic signaling) with rich annotations, based on the peer reviewed literature and existing databases such as CellPhoneDB^10^ and NeuronChatDB^11^; (b) provided several additional tools to allow easy comparisons between multiple datasets/conditions; and (c) added interactive web browser function to allow intuitive exploration and visualization of CellChat outputs. To facilitate intuitive user guided data interpretation, CellChat provides a variety of visualization outputs, including: circle plot, chord diagram, heatmap, hierarchy plot, spatial plot, bubble plot, and cloud plot. Several visualization tools are specially designed for cell-cell communication analysis from spatially resolved transcriptomic data (Fig. 4).

### Applications of the method

To-date, CellChat has been widely used in a broad range of biological systems to dissect signaling mechanisms during tissue homeostasis, development and disease^5^. In our original report^7^, we applied CellChat to scRNA-seq dataset on mouse skin development and predicted a novel role of Edn3 signaling in stimulating directed migration of melanocytes into placodes during mouse hair follicle formation. Comparative analysis of non-lesional and lesional human skin from patients with atopic dermatitis using CellChat uncovered major signaling changes in response to disease. CCL19-CCR7 was identified as the most significant signaling activated in lesional skin, contributing to the communication from inflammatory fibroblasts to dendritic cells. Recently, we used CellChat to study aging-dependent dysregulations during skin wound healing in mice^8^, showing systems level differences in the number, strength, route, and signaling mediators of putative cell-cell communications in young *vs*. aged skin wounds.

Using CellChat, Yang et al. found strong increase in the choroid-to-cortex network across key inflammatory pathways in patients with COVID19 compared to control individuals^12^. Another study revealed increased interactions of CD163/LGMN-macrophages with myofibroblasts, fibroblasts, and pericytes at later time points of COVID19-induced ‘acute respiratory distress syndrome’ (ARDS)^13^. In a single-cell atlas of the adult human cerebral vasculature^14^, CellChat analysis identified Nd2 as the strongest contributor to abnormal cell communications in arteriovenous malformations. Recently, Lake et al. identified state- and niche-dependent signaling for reparative states in proximal and distal tubules using healthy and injured human kidney single-cell atlas^15^. Comparative analysis of *Gabbr1* mutant and control cortices from adult mice uncovered alterations in astrocyte-neuron communication^16^. Ortiz-Muñoz et al. predicted new role for a unique subset of cancer-associated fibroblasts in recruiting monocytes and neutrophils using in situ tumor arrays^17^. A study of PD1 blockade in mismatch repair-deficient colorectal cancer identified interaction between CD4^+^ T helper cells and germinal center B cells in antitumor immunity during immune checkpoint inhibitor treatment^18^.

With the rapid advances in spatially resolved transcriptomics, it is important to characterize cell-cell communication events within a spatial distance for many tissues. For example, in skin, stem cells exclusively located at the bottom of epidermis produce Wnt signals that may only travel short distances to dermis and epidermis^19^. With the spatial location information of cells, CellChat v2 can uncover physically proximal cell-cell communication between interacting cell groups. The added visualization functions in CellChat v2 allows easy exploration of cell-cell communication in the spatial context.

### Comparison with other methods

Numerous computational tools have been developed to facilitate cell-cell communication exploration and analysis^2, 20-28^. The cell-cell communication inference depends on the reference databases of known ligand-receptor interactions. The Python tool CellPhoneDB^22, 29^ is a pioneering method that considers multiple subunits of ligands and receptors to accurately represent known heteromeric molecular complexes. Two other R tools CellChat^7^ and ICELLNET^25^ adopted it, and since then, other tools also followed. Compared to CellPhoneDB and CellChat that have over 2000 ligand-receptor interactions, ICELLNET only has 380 interactions. Recently, CellPhoneDB v4^30^ added interactions of non-protein molecules not directly encoded by genes, and NeuronChat^11^ was designed specifically for neuron-to-neuron communication mediated by neurotransmitters. In CellChat v2, we add new literature-supported interactions, including both proteins and non-proteins acting as ligands, leading to a total of ∼3300 interactions for both mouse and human. Three unique features of CellChat database are: (i) incorporation of soluble and membrane-bound stimulatory and inhibitory cofactors, (ii) classification of ligand-receptor pairs into functionally related signaling pathways, and (iii) rich annotations of each ligand-receptor pair. Feature (i) is considered because many pathways, such as BMP and WNT, are prominently modulated by their cofactors positively or negatively, feature (ii) provides useful insights into signaling mechanisms by examining cell-cell communication at a signaling pathway level, and feature (iii) is useful for selecting ligand-receptor pairs with similar biological functions and interpreting the downstream analysis.

Despite of using different built-in ligand-receptor databases, current tools for inference are all somewhat distinct in their performance, visualization outputs, and downstream analysis. Two recent systematic evaluations of more than 15 cell-cell communication inference methods suggest CellChat is among the top-performing methods^21, 28^. Compared to other tools, CellChat offers a variety of visualization outputs that allow multiple intuitive user-guided interpretations for complex data. One key unique feature of CellChat is its ability to characterize the inferred cell-cell communications using a systems approach. In particular, methods and concepts from social network analysis, pattern recognition, and manifold learning are adapted to derive higher order network information in an easily interpretable way. In addition, CellChat is the pioneering method allowing systematic comparison of communications inferred for different conditions, which is critically important for the ever growing single-cell studies. Later, other methods, such as Connectome^26^, Tensor-cell2cell^27^ and multinichenetr^31^, also introduced functionalities for comparison.

When spatial locations of cells are available, CellChat v2 can identify physically proximal cell-cell communication between interacting cell groups. CellPhoneDB v3 has similar function based on the co-expressed cells within users’ defined microenvironments^23^, while CellChat v2 determines spatially neighboring cell pairs and groups primarily based on an estimated maximal diffusion range of molecular interactions. Both CellChat v2 and CellPhoneDB v3 can eliminate some false positive communication links by utilizing spatial information. Several methods have been specifically designed for spatially resolved transcriptomics, such as SpaOTsc^32^, SpaTalk^33^ and COMMOT^34^. SpaTalk infers cell-cell communication at both single-cell level and cell group level based on whether the ligand-receptor co-expressed cell-cell pairs are nearest neighbors in a KNN graph, using CellTalkDB^35^ as the default database without considering multiple subunits of ligands and receptors. Methods based on KNN graph may neglect communicating cell pairs that are within diffusion ranges, however, not nearest neighbors. SpaOTsc and COMMOT utilized optimal transport methods to quantify the connection between cells with ligands and receptors, with the focus on communication between individual cells. Because of using optimal transport, their computational costs are directly related to the number of cells. Both SpaTalk and COMMOT explore the downstream signaling response, in particular, SpaTalk produces a combined interaction score by integrating inter-cellular and intra-cellular scores.

### Limitations

It is possible that there are missing ligand-receptor interactions not covered in CellChatDB. A tutorial is provided on how to update CellChatDB by adding user-defined ligand-receptor pairs or other resources (Box 1). There are several other limitations for the original CellChat and its v2. First, CellChat infers potential interactions between cell groups without considering heterogeneity within the defined cell groups. To address this limitation, users can refine cell grouping (e.g., spatially heterogenous cells) or define mixed cell types as “super cell types” before applying CellChat v2. Second, similar to other methods, CellChat is limited to hypothesis generation and employs heuristics to guide interpretation of cell-cell communication outputs. With limited benchmarking studies^20, 21, 28^, the question of how to better validate the inferred signaling networks and their downstream gene outputs remains to be answered. Third, cross-condition analysis in CellChat is largely restricted to pairwise comparisons. Identification of signaling changes across multiple conditions and time-series is valuable. Fourth, for molecules that are not directly related to genes measured in scRNA-seq, CellChat v2 estimates the expression of ligands and receptors using those molecules’ key mediators or enzymes for potential communication mediated by non-proteins. Fifth, given that cell-cell communication occurs at the protein level, newly emerging data modalities from single-cell multi-omics, such as single-cell proteomics^36^ and epigenomics^37, 38^, can be used to improve the inference of cell-cell communication. Sixth, CellChat employs a simplified mass action-based model to quantify communication probability between a given ligand and its cognate receptor, and models with more biochemical details can potentially improve inference predictions. Finally, incorporation of the downstream signaling events of activated receptors on receiving cells could further improve the overall inference accuracy^39-43^.

### Expertise needed to implement the protocol

Our protocol requires basic understanding of R and single-cell data analysis, and assumes no specialized bioinformatics training.

### Experimental Design

While CellChat, in principle, can be used for any scRNA-seq and spatially resolved transcriptomic datasets, quality of datasets directly affects the quality of CellChat outputs. First, having sufficient sequencing depth is critical in order to capture gene expression of some (ideally majority of or all) ligands and receptors. For example, expression levels are usually low for ligands during development; thus, sensitivity and depth of sequencing become particularly important for such cases. Second, some image-based spatially resolved transcriptomic techniques, such as MERFISH^44^, need to pre-select genes for measurements. As a result, paired ligand and receptor genes must be measured simultaneously in order to use any cell-cell communication inference methods. Of course, batch effect may introduce output variability for any inference method, including CellChat. Whenever possible, it is important to use the same RNA isolation protocol for replicates and different conditions. More replicates are always helpful for any inference tools to better quantify the reliability of the inference outputs. To perform control analysis, in the protocol we include several datasets that have been well explored using CellChat with known signaling. New CellChat users are encouraged to first test their CellChat code on these datasets by comparing the outputs with the deposited cell-cell communication results.

## Materials

### Hardware

Any desktop workstations or laptops with an Internet connection are sufficient. This protocol was run on a MacBook Pro (MacOS Ventura Monterey, Version 13.5) with 12-Core CPU and 64 GB of RAM. For minimal performance, we recommend using a dual-core CPU with at least 16 GB of RAM on the system for analyses.

#### Software

- Operating system: Linux, Windows (10), or MacOS
- RStudio: an integrated development environment (IDE) for R
- CellChat: the actively maintained open-source program is freely available at https://github.com/jinworks/CellChat.

### Required input data

CellChat requires two user inputs: one is the gene expression data of cells, and the other is the user assigned cell labels. For the gene expression data matrix, genes should be in rows with rownames and cells in columns with colnames. Normalized data is required as input for CellChat analysis, e.g., library-size normalization and then log-transformed with a pseudocount of 1. If user provides raw count data, CellChat provides a `normalizeData` function to do normalization. For the cell group information, a dataframe with rownames is required as input for CellChat. Alternatively, user can take a Seurat, SingleCellExperiment or AnnData object as input. In this case, the meta data in the object is used by default and user needs to provide group.by to define the cell groups. e.g, `group.by = “ident”` for the default cell identities in Seurat object.

When inferring spatially-proximal cell-cell communication from spatially resolved transcriptomic data, user also should provide spatial locations of spots/cells. In addition, to filter out cell-cell communication beyond the maximum diffusion range of molecules (e.g., ∼250 *μm*), CellChat needs to compute the cell-cell distance in the unit of micrometers. Therefore, for spatial technologies like 10x Visium that only provide spatial coordinates in pixels, CellChat converts pixels to micrometers by requiring users to input scale factors and/or spot diameters of the full resolution image. The scale.factors must contain the theoretical spot size (e.g., 65 *μm* for 10x Visium), and the number of pixels that span the diameter of a theoretical spot size in the full-resolution image. The latter can be obtained from the scalefactors_json.json file for 10x Visium.

### Equipment setup

#### Installation of CellChat package

We recommend that users install CellChat and perform analysis in RStudio. In an RStudio environment, the following commands can be run from an R script or directly in the built-in R console. The R commands are the same on MacOS, Linux and Windows.

To install CellChat R packages from GitHub directly, the user should first install the devtools package from the Comprensive R Archive Network (CRAN) and then install CellChat from our GitHub repository by typing in the following commands:

~~~
install.packages(‘devtools’)
devtools::install_github(“jinworks/CellChat”)
~~~

#### Example data

Example datasets for running this protocol can be downloaded from an open access repository figshare at https://figshare.com/projects/Example_data_for_cell-cell_communication_analysis_using_CellChat/157272.

### Procedure

#### Part 1. Inferring cell-cell communication from a single scRNA-seq dataset

▴ CRITICAL The first part of this protocol demonstrates the R commands needed to run the CellChat package for inferring and analyzing cell-cell communication from a single scRNA-seq dataset. These commands can be run from the RStudio integrated development environment or the R command-line interface. The equivalent online version of this described procedure (Steps 1-15) along with the graphical plots are available in the tutorial directory of the CellChat github repository (https://htmlpreview.github.io/?https://github.com/jinworks/CellChat/blob/master/tutorial/CellChat-vignette.html).

#### Data input & preprocessing ●Timing ~12 s

1. Prepare required input data for CellChat analysis. In addition to taking a count data matrix as an input, we also provide instructions for how to prepare CellChat input files from other existing single-cell analysis toolkits, including Seurat and Scanpy. Please start to prepare the input data by following option A when the normalized count data and meta data are available, option B when the Seurat object is available, option C when the SingleCellExperiment object is available, and option D when the Anndata object is available. See details in the online tutorial of the CellChat github repository (https://htmlpreview.github.io/?https://github.com/jinworks/CellChat/blob/master/tutorial/Interface_with_other_single-cell_analysis_toolkits.html).
  (A) **Starting from a count data matrix**

~~~
load(“./tutorial/data_humanSkin_CellChat.rda”)
data.input = data_humanSkin$data # normalized data matrix
meta = data_humanSkin$meta # a dataframe with rownames containing cell mata data
# Subset the input data for CelChat analysis
cell.use = rownames(meta)[meta$condition == “LS”] # extract the cell names from disease data
data.input = data.input[, cell.use]
meta = meta[cell.use,]
~~~
  (B) **Starting from a Seurat object**

~~~
data.input <-GetAssayData(seurat_object, assay = “RNA”, slot = “data”) # normalized data matrix
labels <- Idents(seurat_object)
meta <- data.frame(group = labels, row.names = names(labels)) # c reate a dataframe of the cell labels
~~~
  (C) **Starting from a SingleCellExperiment object**

~~~
data.input <-SingleCellExperiment::logcounts(object) # normalize d data matrix
meta <- as.data.frame(SingleCellExperiment::colData(object)) # ex tract a dataframe of the cell labels
meta$labels <- meta[[“sce.clusters”]]
~~~
  (D) **Starting from an Anndata object**

~~~
# read the data into R using anndata R package
install.packages(“anndata”)
library(anndata)
ad <- read_h5ad(“scanpy_object.h5ad”)
# access count data matrix
counts <- t(as.matrix(ad$X))
# normalize the count data if the normalized data is not available in the .h5ad file
library.size <- Matrix::colSums(counts)
data.input <- as(log1p(Matrix::t(Matrix::t(counts)/library.size) * 10000), “dgCMatrix”)
# access meta data
meta <- ad$obs
meta$labels <- meta[[“clusters”]]
~~~
2. Create a CellChat object by following option A when taking the digital gene expression matrix and cell label information as input, option B when taking a Seurat object as input, option C when taking a SingleCellExperiment object as input, and option D when taking a AnnData object as input. See details in the subsection of **Required input data**. User can initialize a CellChat object using the createCellChat function as follows:
  (A) **Starting from the digital gene expression matrix and cell label information** ▴ CRITICAL If cell mata information is not added when creating the CellChat object, users can also add it later using the `‘addMeta’` function, and set the default cell identities using the `‘setIdent’` function.

~~~
library(CellChat)
cellchat <- createCellChat(object = data.input, meta = meta, group.by = “labels”)
~~~
  (B) **Starting from a Seurat object**

~~~
library(CellChat)
cellChat <- createCellChat(object = seurat.obj, group.by = “ident “, assay = “RNA”)
~~~
  (C) **Starting from a SingleCellExperiment object**

~~~
library(CellChat)
cellChat <- createCellChat(object = sce.obj, group.by = “sce.clusters”)
~~~
  (D) **Starting from an AnnData object**

~~~
sce <- zellkonverter::readH5AD(file = “adata.h5ad”)
# retrieve all the available assays within sce object assayNames(sce)
# add a new assay entry “logcounts” if not available
counts <- assay(sce, “X”) # make sure this is the original count data matrix
library.size <- Matrix::colSums(counts)
logcounts(sce) <- log1p(Matrix::t(Matrix::t(counts)/library.size) * 10000)
# extract a cell meta data
meta <- as.data.frame(SingleCellExperiment::colData(sce)) #
cellChat <- createCellChat(object = sce, group.by = “sce.clusters “)
~~~

♦ TROUBLESHOOTING
3. Before running CellChat to infer cell-cell communication, set the ligand-receptor interaction database and identify over-expressed ligands or receptors. CRITICAL When analyzing human samples, use the database `CellChatDB.human`; when analyzing mouse samples, use the database `CellChatDB.mouse`. CellChatDB includes secreted signaling interactions, extracellular matrix (ECM)-receptor interactions, cell-cell contact interactions as well as interactions involving non-proteins acting as ligands. By default, the interactions involving non-proteins are not used.

~~~
CellChatDB <- CellChatDB.human
showDatabaseCategory(CellChatDB)
# Show the structure of the database
dplyr::glimpse(CellChatDB$interaction)
# use a subset of CellChatDB for cell-cell communication analysis
CellChatDB.use <- subsetDB(CellChatDB, search = “Secreted Signaling”) # use Secreted Signaling
# CellChatDB.use <- CellChatDB # simply use the default CellChatDB to use all CellChatDB for cell-cell communication analysis
# set the used database in the object
cellchat@DB <- CellChatDB.use
~~~
4. Identify over-expressed ligands or receptors. To infer the cell state-specific communications, CellChat identifies over-expressed ligands or receptors in one cell group and then identifies over-expressed ligand-receptor interactions if either ligand or receptor are over-expressed.

~~~
cellchat <- subsetData(cellchat) # This step is necessary even if using the whole database
future::plan(“multisession”, workers = 4) # do parallel
cellchat <- identifyOverExpressedGenes(cellchat)
cellchat <- identifyOverExpressedInteractions(cellchat)
~~~

♦ TROUBLESHOOTING

### Inference of cell-cell communication network ● Timing ~37 s

5. Infer cell-cell communication at a ligand-receptor pair level. Users are now ready to infer cell-cell communication by using the following command:

~~~
cellchat <- computeCommunProb(cellchat, type = “triMean”)
~~~ ▴ CRITICAL The key parameter for this analysis is `type`, the method for computing the average gene expression per cell group. By default `type` = “triMean”, producing fewer but stronger interactions. When setting ‘type = “truncatedMean”‘, a value should be assigned to ‘trim’, producing more interactions. By default, CellChat uses a statistically robust mean method called ‘trimean’, which produces fewer interactions than other methods. However, we find that CellChat performs well at predicting stronger interactions, which is very helpful for narrowing down on interactions for further experimental validations. The ‘trimean’ approximates 25% truncated mean, implying that the average gene expression is zero if the percent of expressed cells in one group is less than 25%. To identify weak signaling, users should use “truncatedMean”. In general, users can use 10% truncated mean by setting ‘type = “truncatedMean”’ and ‘trim = 0.1’. When using “triMean”, “truncatedMean” with ‘trim = 0.1’, and “truncatedMean” with ‘trim = 0.05’, Figs. 3a and 3b show an increase in the number of inferred interactions and in the enriched signaling pathways respectively. To determine a proper value of `trim`, CellChat provides a function ‘`computeAveExpr`’, which can help to check the average expression of signaling genes of interest, e.g, ‘`computeAveExpr(cellchat, features = c(“CXCL12”,”CXCR4”), type = “truncatedMean”, trim = 0.1)`’. Therefore, if well-known signaling pathways in the studied biological process are not predicted, users can try ‘truncatedMean’ with lower values of ‘trim’ to change the method for calculating the average gene expression per cell group. ♦ TROUBLESHOOTING ▴ CAUTION Another important parameter is raw.use. It indicates whether to use the raw data (i.e., ‘object@data.signaling’) or the projected data (i.e., ‘object@data.project’). The default is TRUE. Set raw.use = FALSE to use the projected data when analyzing single-cell data with shallow sequencing depth because the projected data could help to reduce the dropout effects of signaling genes, in particular for possible zero expression of subunits of ligands/receptors. To use the projected data, users should run the function `projectData` before running `computeCommunProb`. The function `projectData` smooths genes’ expression values based on their neighbors’ defined in a high-confidence experimentally validated protein-protein network. When using the projected data, the number of inferred interactions clearly increases. However, generally, it only introduces very weak communications.
6. Filter the cell-cell communication. Users can filter out the cell-cell communication if there are only few cells in certain cell groups. By default, the minimum number of cells required in each cell group for cell-cell communication is 10.

~~~
cellchat <- filterCommunication(cellchat, min.cells = 10)
~~~
7. Infer cell-cell communication at a signaling pathway level. CellChat computes the communication probability on signaling pathway level by summarizing the communication probabilities of all ligands-receptors interactions associated with each signaling pathway. Note that the inferred intercellular communication network of each ligand-receptor pair and each signaling pathway is stored in the slot ‘net’ and ‘netP’, respectively.

~~~
cellchat <- computeCommunProbPathway(cellchat)
~~~
8. Calculate aggregated cell-cell communication network. CellChat can calculate the aggregated cell-cell communication network by counting the number of links or summarizing the communication probability. Users can also calculate the aggregated network among a subset of cell groups by setting ‘sources.use’ and ‘targets.use’.

~~~
cellchat <- aggregateNet(cellchat)
~~~

◼ PAUSE POINT Users can export the CellChat object together with the inferred cell-cell communication networks and save them for later use:

~~~
saveRDS(cellchat, file = “cellchat_humanSkin_LS.rds”)
~~~

#### Visualization of cell-cell communication network ● Timing ~10 s

▴ CRITICAL Upon inferring the cell-cell communication network, CellChat provides various functionalities for further data exploration, analysis, and visualization. Specifically:

- It provides several ways for visualizing cell-cell communication network, including hierarchical plot, circle plot, chord diagram, and bubble plot.
- It provides an easy-to-use tool for extracting and visualizing high-order information of the inferred networks. For example, it allows prediction of major signaling inputs and outputs for cell populations and how these populations and signals coordinate together for functions.
- It can quantitatively characterize and compare the inferred cell-cell communication networks using an integrated approach by combining concepts from social network analysis, pattern recognition, and manifold learning approaches.
  9. Visualize each signaling pathway using Circle plot (option A), Hierarchy plot (option B), Chord diagram (option C) and Heatmap (option D). Users can visualize the inferred communication network of signaling pathways using ‘netVisual_aggregate’, and visualize the inferred communication networks of individual L-R pairs associated with that signaling pathway using ‘netVisual_individual’. In all visualization plots, edge colors are consistent with the sources as sender, and edge weights are proportional to the interaction strength. Thicker edge line indicates a stronger signal. All the signaling pathways showing significant communications can be accessed by ‘cellchat@netP$pathways’. Here we take input of one signaling pathway as an example.
    (A) **Circle plot**

~~~
# Access all the signaling pathways showing significant communications
pathways.show.all <- cellchat@netP$pathways
# select one pathway
pathways.show <- c(“CXCL”)
par(mfrow=c(1,1))
netVisual_aggregate(cellchat, signaling = pathways.show, layout = “circle”, color.use = NULL, sources.use = NULL, targets.use = NULL, idents.use = NULL)
~~~ ♦ TROUBLESHOOTING
    (B) **Hierarchy plot**

~~~
vertex.receiver = seq(1,4)
netVisual_aggregate(cellchat, signaling = pathways.show, layout = “hierarchy”, vertex.receiver = vertex.receiver)
~~~ ▴ CRITICAL The key parameter for this plot is `vertex.receiver`, a numeric vector giving the index of the cell groups as targets in the left part of hierarchy plot. This hierarchical plot consist of two components: the left portion shows autocrine and paracrine signaling to certain cell groups of interest (i.e, the defined ‘vertex.receiver’), and the right portion shows autocrine and paracrine signaling to the remaining cell groups in the dataset. Thus, hierarchy plot provides an informative and intuitive way to visualize autocrine and paracrine signaling communications between cell groups of interest. In the above example, to study the cell-cell communication between fibroblasts and immune cells, we define ‘vertex.receiver’ as all fibroblast cell groups. In the hierarchy plot, solid and open circles represent source and target, respectively.
    (C) **Chord diagram**
      i. Create a chord diagram using the universal function ‘`netVisual_aggregate’`. In the chord diagram, the inner thinner bar colors represent the targets that receive signal from the corresponding outer bar. The inner bar size is proportional to the signal strength received by the targets. Such inner bar is helpful for interpreting the complex chord diagram.

~~~
par(mfrow=c(1,1))
netVisual_aggregate(cellchat, signaling = pathways.show, layout = “chord”)
~~~
      ii. Customize a chord diagram using two independent functions ’`netVisual_chord_cell`’ and ‘`netVisual_chord_gene`’ to flexibly visualize cell-cell communication with different purposes and at different levels. ‘netVisual_chord_cell’ is used for visualizing the cell-cell communication between different cell groups (where each sector in the chord diagram is a cell group), and ‘netVisual_chord_gene’ is used for visualizing the cell-cell communication mediated by multiple ligand-receptors or signaling pathways (where each sector in the chord diagram is a ligand, receptor or signaling pathway). These two functions can flexibly visualize the signaling network by adjusting different parameters in the circlize (https://github.com/jokergoo/circlize) package. For example, we can define a named char vector ‘group’ to create multiple-group chord diagram, e.g., grouping cell clusters into different cell types.

~~~
par(mfrow=c(1,1))
group.cellType <- c(rep(“FIB”, 4), rep(“DC”, 4), rep(“TC”, 4)) # grouping cell clusters into fibroblast, DC and TC cells
names(group.cellType) <- levels(cellchat@idents)
netVisual_chord_cell(cellchat, signaling = pathways.show, group = group.cellType, title.name = paste0(pathways.show, “signaling n etwork”))
~~~
    (D) **Heatmap plot**

~~~
par(mfrow=c(1,1))
netVisual_heatmap(cellchat, signaling = pathways.show, color.heat map = “Reds”)
~~~
  10. Visualize inferred communication networks of individual L-R pairs. CellChat first can compute the contribution of each ligand-receptor pair with one particular signaling pathway, and then visualize the cell-cell communication mediated by a single ligand-receptor pair. CellChat provides a function ‘`extractEnrichedLR`’ to extract all the significant interactions (L-R pairs) and related signaling genes for a given signaling pathway:

~~~
netAnalysis_contribution(cellchat, signaling = pathways.show)
pairLR.CXCL <- extractEnrichedLR(cellchat, signaling = pathways.s how, geneLR.return = FALSE)
LR.show <- pairLR.CXCL[1,] # show one ligand-receptor pair
netVisual_individual(cellchat, signaling = pathways.show, pairLR. use = LR.show, layout = “circle”)
~~~
  11. Visualize cell-cell communication mediated by multiple ligand-receptors or signaling pathways. CellChat can also show all the significant interactions (L-R pairs), interactions mediated by certain signaling pathways as well as interactions provided by users from some cell groups to other cell groups using the function `netVisual_bubble (`option A) and `netVisual_chord_gene` (option B).
    (A) **Bubble plot**

~~~
# (1) show all the significant interactions (L-R pairs) from some
 cell groups (defined by ‘sources.use’) to other cell groups (defined by ‘targets.use’)
netVisual_bubble(cellchat, sources.use = 4, targets.use = c(5:1 1), remove.isolate = FALSE)
# (2) show all the significant interactions (L-R pairs) associated with certain signaling pathways
netVisual_bubble(cellchat, sources.use = 4, targets.use = c(5:1 1), signaling = c(“CCL”,”CXCL”), remove.isolate = FALSE)
# (3) show all the significant interactions (L-R pairs) based on user’s input (defined by ‘pairLR.use’)
pairLR.use <- extractEnrichedLR(cellchat, signaling = c(“CCL”,”CX CL”,”FGF”))
netVisual_bubble(cellchat, sources.use = c(3,4), targets.use = c (5:8), pairLR.use = pairLR.use, remove.isolate = TRUE)
# set the order of interacting cell pairs on x-axis
# (4) Default: first sort cell pairs based on the appearance of sources in levels(object@idents), and then based on the appearance of targets in levels(object@idents)
# (5) sort cell pairs based on the targets.use defined by users
netVisual_bubble(cellchat, targets.use = c(“LC”,”Inflam. DC”,”cDC 2”,”CD40LG+ TC”), pairLR.use = pairLR.use, remove.isolate = TRUE, sort.by.target = T)
# (6) sort cell pairs based on the sources.use defined by users
netVisual_bubble(cellchat, sources.use = c(“FBN1+ FIB”,”APOE+ FIB “,”Inflam. FIB”), pairLR.use = pairLR.use, remove.isolate = TRUE, sort.by.source = T)
# (7) sort cell pairs based on the sources.use and then targets.use defined by users
netVisual_bubble(cellchat, sources.use = c(“FBN1+ FIB”,”APOE+ FIB “,”Inflam. FIB”), targets.use = c(“LC”,”Inflam. DC”,”cDC2”,”CD40L G+ TC”), pairLR.use = pairLR.use, remove.isolate = TRUE, sort.by. source = T, sort.by.target = T)
# (8) sort cell pairs based on the targets.use and then sources.u se defined by users
netVisual_bubble(cellchat, sources.use = c(“FBN1+ FIB”,”APOE+ FIB “,”Inflam. FIB”), targets.use = c(“LC”,”Inflam. DC”,”cDC2”,”CD40L G+ TC”), pairLR.use = pairLR.use, remove.isolate = TRUE, sort.by. source = T, sort.by.target = T, sort.by.source.priority = FALSE)
~~~ ▴ CRITICAL Important parameters of the netVisual_bubble function are as follows:
      - slot.name: the slot name of object: slot.name = “net” when visualizing links at the level of ligands/receptors; slot.name = “netP” when visualizing links at the level of signaling pathways;
      - targets.use: a vector giving the index or the name of target cell groups;
      - signaling: a character vector giving the name of signaling pathways of interest;
      - pairLR.use: a data frame consisting of one column named either “interaction_name” or “pathway_name”, defining the interactions of interest and the order of L-R on y-axis;
      - remove.isolate: whether to remove the entire empty column, i.e., communication between certain cell groups.
      - sort.by.source, sort.by.target, sort.by.source.priority: reorder the interacting cell pairs
    (B) **Chord diagram**

~~~
# show all the significant interactions (L-R pairs) from some cell groups (defined by ‘sources.use’) to other cell groups (defined by ‘targets.use’)
# show all the interactions sending from Inflam.FIB
netVisual_chord_gene(cellchat, sources.use = 4, targets.use = c (5:11), lab.cex = 0.5,legend.pos.y = 30)
# show all the interactions received by Inflam.DC
netVisual_chord_gene(cellchat, sources.use = c(1,2,3,4), targets. use = 8, legend.pos.x = 15)
# show all the significant interactions (L-R pairs) associated with certain signaling pathways
netVisual_chord_gene(cellchat, sources.use = c(1,2,3,4), targets. use = c(5:11), signaling = c(“CCL”,”CXCL”),legend.pos.x = 8)
# show all the significant signaling pathways from some cell groups (defined by ‘sources.use’) to other cell groups (defined by ‘t argets.use’)
netVisual_chord_gene(cellchat, sources.use = c(1,2,3,4), targets. use = c(5:11), slot.name = “netP”, legend.pos.x = 10)
~~~ ▴ CRITICAL Important parameters of the `netVisual_chord_gene` function are as follows:
      - slot.name: the slot name of object: slot.name = “net” when visualizing links at the level of ligands/receptors; slot.name = “netP” when visualizing links at the level of signaling pathways;
      - signaling: a character vector giving the name of signaling networks;
      - pairLR.use: a data frame consisting of one column named either “interaction_name” or “pathway_name”, defining the interactions of interest;
      - net: a data frame consisting of the interactions of interest. “net” needs to have at least three columns: “source”,”target” and “interaction_name” when visualizing links at the level of ligands/receptors; “source”,”target” and “pathway_name” when visualizing links at the level of signaling pathway; “interaction_name” and “pathway_name” must be the matched names in “CellChatDB$interaction”;
      - sources.use: a vector giving the index or the name of source cell groups;
      - targets.use: a vector giving the index or the name of target cell groups;
      - color.use: colors for the cell groups;
      - lab.cex: font size for the text;
      - small.gap: small gap between sectors; If the gene names are overlapping, users can adjust the argument ‘small.gap’ by decreasing their values.
      - big.gap: gap between the different sets of sectors, which are defined in the ‘group’ parameter;
  12. Plot signaling gene expression distribution. CellChat can plot the gene expression distribution of signaling genes related to L-R pairs or signaling pathways using a Seurat wrapper function ‘`plotGeneExpression`’ if the Seurat R package has been installed. Alternatively, users can extract the signaling genes related to the inferred L-R pairs or signaling pathway using ‘extractEnrichedLR’, and then plot gene expression using Seurat or other packages.

~~~
plotGeneExpression(cellchat, signaling = “CXCL”, enriched.only = TRUE)
~~~

#### Systems analysis of cell-cell communication network ● Timing ~ 2.7 mins

▴ CRITICAL To facilitate the interpretation of the complex intercellular communication networks, CellChat quantitively measures networks through methods abstracted from graph theory, pattern recognition and manifold learning.

- It can determine major signaling sources and targets as well as mediators and influencers within a given signaling network using centrality measures from network analysis;
- It can predict key incoming and outgoing signals for specific cell types as well as coordinated responses among different cell types by leveraging pattern recognition approaches;
- It can group signaling pathways by defining similarity measures and performing manifold learning from both functional and topological perspectives.
  13. Identify signaling roles of cell groups as well as the major contributing signaling. CellChat allows identification of dominant senders, receivers, mediators and influencers in the intercellular communication network by computing several network centrality measures for each cell group. Specifically, CellChat uses measures in weighted-directed networks, including out-degree, in-degree, flow betweenness and information centrality^7, 45^, to respectively identify dominant senders, receivers, mediators and influencers for the intercellular communications. In a weighted directed network with the weights as the computed communication probabilities, the outdegree, computed as the sum of communication probabilities of the outgoing signaling from a cell group, and the in-degree, computed as the sum of the communication probabilities of the incoming signaling to a cell group, can be used to identify the dominant cell senders and receivers of signaling networks, respectively. Users can visualize the centrality scores on a heatmap (option A) and a 2D plot (option B). CellChat can also answer the question on which signals contribute the most to outgoing or incoming signaling of certain cell groups (option C).
    (A) **Compute and visualize the network centrality scores**

~~~
# Compute the network centrality scores
cellchat <- netAnalysis_computeCentrality(cellchat, slot.name = “netP”) # the slot ‘netP’ means the inferred intercellular communication network of signaling pathways
# Visualize the computed centrality scores using heatmap, allowing ready identification of major signaling roles of cell groups
netAnalysis_signalingRole_network(cellchat, signaling = pathways. show, width = 8, height = 2.5, font.size = 10)
~~~ ♦TROUBLESHOOTING
    (B) **Visualize dominant senders (sources) and receivers (targets) in a 2D space**

~~~
netAnalysis_signalingRole_scatter(cellchat, signaling = NULL)
~~~ ▴ CRITICAL CellChat also provides another intuitive way to visualize the dominant senders (sources) and receivers (targets) in a 2D space using scatter plot. This scatter plot shows the dominant senders (sources) and receivers (targets) in a 2D space. X-axis and y-axis are respectively the total outgoing or incoming communication probability associated with each cell group. Dot size is proportional to the number of inferred links (both outgoing and incoming) associated with each cell group. Dot colors indicate different cell groups. Dot shapes indicate different categories of cell groups if ‘group’’ is defined. Important parameters of this function are as follows:
      - signaling: a char vector to specify signaling pathway names of interest. signaling = NULL: Signaling role analysis on the aggregated cell-cell communication network from all signaling pathways;
      - color.use: defining the color for each cell group;
      - slot.name: the slot name of object that is used to compute centrality measures of signaling networks;
      - group: a vector to categorize the cell groups, e.g., categorize the cell groups into two major categories: immune cells and fibroblasts;
      - x.measure: The measure used as x-axis. This measure should be one of ‘names(slot(object, slot.name)$centr[[1]])’ computed from ‘netAnalysis_computeCentrality’. Default = “outdeg” is the weighted outgoing links (i.e., Outgoing interaction strength). If setting as “outdeg_unweighted”, it represents the total number of outgoing signaling;
      - y.measure: The measure used as y-axis. This measure should be one o ‘names(slot(object, slot.name)$centr[[1]])’ computed from ‘netAnalysis_computeCentrality’. Default = “indeg” is the weighted incoming links (i.e., Incoming interaction strength). If setting as “indeg_unweighted”, it represents the total number of incoming signaling.
    (C) **Identify signals contributing the most to outgoing or incoming signaling of certain cell groups**

~~~
ht1 <- netAnalysis_signalingRole_heatmap(cellchat, pattern = “out going”)
ht2 <- netAnalysis_signalingRole_heatmap(cellchat, pattern = “incoming”)
ht1 + ht2
~~~ ▴ CRITICAL In this heatmap, colobar represents the relative signaling strength of a signaling pathway across cell groups (NB: values are row-scaled). The top colored bar plot shows the total signaling strength of a cell group by summarizing all signaling pathways displayed in the heatmap. The right grey bar plot shows the total signaling strength of a signaling pathway by summarizing all cell groups displayed in the heatmap. Important parameters of this function are as follows:
      - signaling: a character vector giving the name of signaling networks;
      - pattern: “outgoing”, “incoming” or “all”. When pattern = “all”, it aggregates the outgoing and incoming signaling strength together;
      - slot.name: the slot name of object that is used to compute centrality measures of signaling networks;
      - color.use: the character vector defining the color of each cell group;
  14. Identify global communication patterns to explore how multiple cell types and signaling pathways coordinate together. In addition to exploring detailed communications for individual pathways, an important question is how multiple cell groups and signaling pathways coordinate to function. CellChat employs a pattern recognition method to identify the global communication patterns. As the number of patterns increases, there might be redundant patterns, making it difficult to interpret the communication patterns. We chose five patterns as default. Generally, it is biologically meaningful with the number of patterns greater than two. In addition, CellChat also provides a function ‘`selectK`’ to infer the number of patterns, which is based on two metrics that have been implemented in the NMF R package, including Cophenetic and Silhouette. Both metrics measure the stability for a particular number of patterns based on a hierarchical clustering of the consensus matrix. For a range of the number of patterns, a suitable number of patterns is the one at which Cophenetic and Silhouette values begin to drop suddenly. This analysis can be done for outgoing (option A) and incoming (option B) signaling patterns. Outgoing patterns reveal how the sender cells (i.e., cells as signal source) coordinate with each other as well as how they coordinate with certain signaling pathways to drive communication. Incoming patterns show how the target cells (i.e., cells as signal receivers) coordinate with each other as well as how they coordinate with certain signaling pathways to respond to incoming signals.
    (A) **Identify and visualize outgoing communication pattern of secreting cells**

~~~
# infer the number of patterns.
selectK(cellchat, pattern =“outgoing”)
nPatterns = 3 # Both Cophenetic and Silhouette values begin to drop suddenly when the number of outgoing patterns is 3.
cellchat <- identifyCommunicationPatterns(cellchat, pattern =“out going”, k = nPatterns)
# river plot
netAnalysis_river(cellchat, pattern =“outgoing”)
# dot plot
netAnalysis_dot(cellchat, pattern =“outgoing”)
~~~ CRITICAL To intuitively show the associations of latent patterns with cell groups and ligand-receptor pairs or signaling pathways, we used a river (alluvial) plot. We first normalized each row of W and each column of H to be [0,1], and then set the elements in W and H to be zero if they are less than 0.5. Such thresholding allows to uncover the most enriched cell groups and signaling pathways associated with each inferred pattern, that is, each cell group or signaling pathway is associated with only one inferred pattern. These thresholded matrices W and H are used as inputs for creating alluvial plots. To directly relate cell groups with their enriched signaling pathways, we set the elements in W and H to be zero if they are less than 1/R where R is the number of latent patterns. By using a less strict threshold, more enriched signaling pathways associated each cell group might be obtained. Using a contribution score of each cell group to each signaling pathway computed by multiplying W by H, we constructed a dot plot in which the dot size is proportion to the contribution score to show association between cell group and their enriched signaling pathways. Users can also decrease the parameter ‘cutoff’ to show more enriched signaling pathways associated each cell group.
    (B) **Identify and visualize incoming communication pattern of target cells**

~~~
selectK(cellchat, pattern = “incoming”)
nPatterns = 4
cellchat <- identifyCommunicationPatterns(cellchat, pattern = “incoming”, k = nPatterns)
# river plot
netAnalysis_river(cellchat, pattern = “incoming”)
# dot plot
netAnalysis_dot(cellchat, pattern = “incoming”)
~~~
  15. Manifold and classification learning analysis of signaling networks ▴ CRITICAL CellChat is able to quantify the similarity between all significant signaling pathways and then group them based on their cellular communication network similarity. This analysis is helpful to predict putative functions of the poorly studied pathways based on their similarity with the pathways with well-known functions. Grouping can be done either based on the functional (following option A) or structural similarity (following option B). In terms of functional similarity, high degree of functional similarity indicates major senders and receivers are similar, and it can be interpreted as the two signaling pathways or two ligand-receptor pairs exhibit similar and/or redundant roles. In terms of structural similarity, a structural similarity was used to compare their signaling network structure, without considering the similarity of senders and receivers. To obtain a manifold embedding of all inferred communication networks, we first compute the pairwise functional or topological similarity between any pair of inferred networks, then smooth the similarity matrix using a shared nearest neighbor (SNN) graph and finally perform UMAP on the smoothed similarity matrix. It should be noted that functional similarity analysis is not applicable to multiple datasets with different cell type composition, but structural similarity analysis is applicable to multiple datasets either with the same cell type composition or with the vastly different cell type composition. More detailed information is available at our previous study^7^.
    (A) **Functional similarity**

~~~
cellchat <- computeNetSimilarity(cellchat, type = “functional”)
cellchat <- netEmbedding(cellchat, type = “functional”)
cellchat <- netClustering(cellchat, type = “functional”)
# Visualization in 2D-space
netVisual_embedding(cellchat, type = “functional”, label.size = 3.5)
# netVisual_embeddingZoomIn(cellchat, type = “functional”, nCol = 2)
~~~
    (B) **Structure similarity**

~~~
cellchat <- computeNetSimilarity(cellchat, type = “structural”)
cellchat <- netEmbedding(cellchat, type = “structural”)
cellchat <- netClustering(cellchat, type = “structural”)
# Visualization in 2D-space
netVisual_embedding(cellchat, type = “structural”, label.size = 3.5)
# netVisual_embeddingZoomIn(cellchat, type = “structural”, nCol = 2)
~~~

◼PAUSE POINT Users can now export the CellChat object and save them for later use:

~~~
saveRDS(cellchat, file = “cellchat_humanSkin_LS.rds”)
~~~

#### Part 2. Comparative analysis of cell-cell communication from pairs of scRNA-seq datasets

▴ CRITICAL In this section, we demonstrate CellChat’s ability to identify major signaling changes across different biological conditions by quantitative contrasts and joint manifold learning. We showcase CellChat’s diverse functionalities by applying it to a scRNA-seq data on cells from two biological conditions: nonlesional (NL, normal) and lesional (LS, diseased) human skin from patients with atopic dermatitis. These two datasets (conditions) have the same cell population compositions after joint clustering. If there are different cell population compositions between different conditions, please check out the procedure of Part 3. The equivalent online version of this described procedure (Steps 16-26) along with the graphical plots are available in the tutorial directory of the CellChat github repository (https://htmlpreview.github.io/?https://github.com/jinworks/CellChat/blob/master/tutorial/Comparison_analysis_of_multiple_datasets.html).

#### Load CellChat object of each dataset and merge them together ● Timing ~3 s

16. Merge multiple CellChat objects for comparison analysis. Users need to first run CellChat on each dataset separately (Steps 1-8) and then merge different CellChat objects together. Please do ‘updateCellChat’ if the CellChat objects are obtained using the earlier version (< 1.6.0).

~~~
library(CellChat)
library(patchwork)
cellchat.NL <- readRDS(“./tutorial/cellchat_humanSkin_NL.rds”)
cellchat.LS <- readRDS(“./tutorial/cellchat_humanSkin_LS.rds”)
cellchat.NL <- updateCellChat(cellchat.NL)
cellchat.LS <- updateCellChat(cellchat.LS)
object.list <- list(NL = cellchat.NL, LS = cellchat.LS)
cellchat <- mergeCellChat(object.list, add.names = names(object.list))
~~~

◼PAUSE POINT Users can now export the merged CellChat object and the list of the two separate objects for later use:

~~~
save(object.list, file = “cellchat_object.list_humanSkin_NL_LS.RData”)
save(cellchat, file = “cellchat_merged_humanSkin_NL_LS.RData”)
~~~

#### Predict general principles of cell-cell communication ● Timing ~2 s

CRITICAL CellChat employs a top-down approach, i.e., starting with the big picture and then refining it in a greater detail on the signaling mechanisms, to identify signaling changes at different levels, including both general principles of cell-cell communication and dysfunctional cell populations, signaling pathways and ligand-receptors. First, CellChat starts with a big picture to predict general principles of cell-cell communication. When comparing cell-cell communication among multiple biological conditions, it can answer the following biological questions:

- Whether the cell-cell communication is enhanced or not;
- The interaction between which cell types is significantly changed;
- How the major sources and targets change from one condition to another.
  17. Compare the total number of interactions and interaction strength. To answer the question on whether the cell-cell communication is enhanced or not, CellChat compares the total number of interactions and interaction strength of the inferred cell-cell communication networks from different biological conditions.

~~~
gg1 <- compareInteractions(cellchat, show.legend = F, group = c (1,2))
gg2 <- compareInteractions(cellchat, show.legend = F, group = c(1,2), measure = “weight”)
gg1 + gg2
~~~
  18. Compare the number of interactions and interaction strength among different cell populations. To identify the interaction between cell populations showing significant changes, CellChat compares the number of interactions and interaction strength among different cell populations using a circle plot with differential interactions (option A), a heatmap with differential interactions (option B) and two circle plots with the number of interactions or interaction strength per dataset (option C). Alternatively, users can examine the differential number of interactions or interaction strength among coarse cell types by aggregating the cell-cell communication based on the defined cell groups (option D).
    (A) **Circle plot showing differential number of interactions or interaction strength among different cell populations across two datasets** CRITICAL The differential number of interactions or interaction strength in the cell-cell communication network between two datasets can be visualized using circle plot, where red (or blue) colored edges represent increased (or decreased) signaling in the second dataset compared to the first one.

~~~
par(mfrow = c(1,2), xpd=TRUE)
netVisual_diffInteraction(cellchat, weight.scale = T)
netVisual_diffInteraction(cellchat, weight.scale = T, measure = “weight”)
~~~ ♦TROUBLESHOOTING
    (B) **Heatmap showing differential number of interactions or interaction strength among different cell populations across two datasets** CRITICAL CellChat can also show differential number of interactions or interaction strength in greater details using a heatmap. The top colored bar plot represents the sum of column of values displayed in the heatmap (incoming signaling). The right colored bar plot represents the sum of row of values (outgoing signaling). In the colorbar, red (or blue) colored edges represent increased (or decreased) signaling in the second dataset compared to the first one.

~~~
gg1 <- netVisual_heatmap(cellchat)
gg2 <- netVisual_heatmap(cellchat, measure = “weight”)
gg1 + gg2
~~~
    (C) **Circle plot showing the number of interactions or interaction strength among different cell populations across multiple datasets** CRITICAL The differential network analysis only works for pairwise datasets. If there are more datasets for comparison, CellChat can directly show the number of interactions or interaction strength between any two cell populations in each dataset. To better control the node size and edge weights of the inferred networks across different datasets, CellChat computes the maximum number of cells per cell group and the maximum number of interactions (or interaction weights) across all datasets.

~~~
weight.max <- getMaxWeight(object.list, attribute = c(“idents”,”count”))
par(mfrow = c(1,2), xpd=TRUE)
for (i in 1:length(object.list)) {
 netVisual_circle(object.list[[i]]@net$count, weight.scale = T, label.edge= F, edge.weight.max = weight.max[2], edge.width.max = 12, title.name = paste0(“Number of interactions - “, names(object.list)[i]))
}
~~~
    (D) **Circle plot showing the differential number of interactions or interaction strength among coarse cell types** CRITICAL To simplify the complicated network and gain insights into the cell-cell communication at the cell type level, CellChat can aggregate the cell-cell communication based on the defined cell groups.

~~~
# Here, CellChat categorize the cell populations into three cell types, and then re-merge the list of CellChat objects.
group.cellType <- c(rep(“FIB”, 4), rep(“DC”, 4), rep(“TC”, 4))
group.cellType <- factor(group.cellType, levels = c(“FIB”, “DC”, “TC”))
object.list <- lapply(object.list, function(x) {mergeInteractions (x, group.cellType)})
cellchat <- mergeCellChat(object.list, add.names = names(object.l ist))
# Show the number of interactions or interaction strength between any two cell types in each dataset.
weight.max <- getMaxWeight(object.list, slot.name = c(“idents”, “net”, “net”), attribute = c(“idents”,”count”, “count.merged”))
par(mfrow = c(1,2), xpd=TRUE)
for (i in 1:length(object.list)) {
 netVisual_circle(object.list[[i]]@net$count.merged, weight.scal e = T, label.edge= T, edge.weight.max = weight.max[3], edge.width.max = 12, title.name = paste0(“Number of interactions - “, name s(object.list)[i]))
}
# Similarly, CellChat can also show the differential number of in teractions or interaction strength between any two cell types using circle plot.
par(mfrow = c(1,2), xpd=TRUE)
netVisual_diffInteraction(cellchat, weight.scale = T, measure = “count.merged”, label.edge = T)
netVisual_diffInteraction(cellchat, weight.scale = T, measure = “weight.merged”, label.edge = T)
~~~
  19. Compare major sources and targets in 2D space. Identify cell populations with significant changes in sending or receiving signals between different datasets by following option A, and the signaling changes of specific cell populations by following option B.
    (A) **Identify cell populations with significant changes in sending or receiving signals**

~~~
num.link <- sapply(object.list, function(x) {rowSums(x@net$count) + colSums(x@net$count)-diag(x@net$count)})
weight.MinMax <- c(min(num.link), max(num.link)) # control the dot size in the different datasets
gg <- list()
for (i in 1:length(object.list)) {
 gg[[i]] <- netAnalysis_signalingRole_scatter(object.list[[i]], title = names(object.list)[i], weight.MinMax = weight.MinMax)
}
patchwork::wrap_plots(plots = gg)
~~~ ♦TROUBLESHOOTING
    (B) **Identify the signaling changes of specific cell populations**

~~~
gg1 <- netAnalysis_signalingChanges_scatter(cellchat, idents.use = “Inflam. DC”, signaling.exclude = “MIF”)
gg2 <- netAnalysis_signalingChanges_scatter(cellchat, idents.use = “cDC1”, signaling.exclude = c(“MIF”))
patchwork::wrap_plots(plots = list(gg1,gg2))
~~~

#### Identify altered signaling with distinct network architecture and interaction strength ● Timing ~17 s

20. Identify signaling networks with larger (or smaller) difference as well as signaling groups based on their functional/structure similarity. CellChat performs joint manifold learning and classification of all inferred communication networks across different conditions. The manifold embeddings are obtained by first computing the pairwise functional or topological similarity between any pair of inferred networks, and then performing UMAP on a SNN-smoothed similarity matrix. More detailed information of the functional and structural similarity is described in Step 15. By quantifying the similarity between the cellular communication networks of signaling pathways across conditions, this analysis highlights the potentially altered signaling pathway. CellChat adopts the concept of network rewiring from network biology and hypothesized that the difference between different communication networks may affect biological processes across conditions. UMAP is used for visualizing signaling relationship and interpreting our signaling outputs in an intuitive way without involving the classification of conditions.

~~~
cellchat <- computeNetSimilarity(cellchat, type = “functional”)
cellchat <- netEmbedding(cellchat, type = “functional”)
cellchat <- netClustering(cellchat, type = “functional”)
# Visualization in 2D-space
netVisual_embedding(cellchat, type = “functional”, label.size = 3.5)
netVisual_embeddingZoomIn(cellchat, type = “functional”, nCol = 2)
# Compute and visualize the pathway distance in the learned joint manifold
rankSimilarity(cellchat, type = “functional”)
~~~ CRITICAL CellChat can identify the signaling networks with larger (or smaller) difference based on their Euclidean distance in the shared two-dimensions space. Larger distance implies larger difference of the communication networks between two datasets in terms of either functional or structure similarity. It should be noted that we only compute the distance of overlapped signaling pathways between two datasets. Those signaling pathways that are only identified in one dataset are not considered here. If there are more than three datasets, one can do pairwise comparisons by modifying the parameter ‘comparison’ in the function ‘`rankSimilarity`’.
21. Identify altered signaling with distinct interaction strength. By comparing the information flow/interaction strength of each signaling pathway, CellChat can identify signaling pathways that: (i) turn off, (ii) decrease, (iii) turn on, or (iv) increase, by changing their information flow at one condition as compared to another condition. Identify the altered signaling pathways based on the overall information flow by following option A, and identify signaling pathways/ligand-receptors that exhibit different signaling patterns by following option B.
  (A) **Compare the overall information flow of each signaling pathway** CRITICAL CellChat can identify the altered signaling pathways by simply comparing the information flow for each signaling pathway, which is defined by the sum of communication probability among all pairs of cell groups in the inferred network (i.e., the total weights in the network). This bar graph can be plotted in a stacked mode or not. Significant signaling pathways are ranked based on differences in the overall information flow within the inferred networks between NL and LS skin. The top signaling pathways colored red are enriched in NL skin, and these colored greens are enriched in the LS skin.

~~~
gg1 <- rankNet(cellchat, mode = “comparison”, stacked = T, do.stat = TRUE)
gg2 <- rankNet(cellchat, mode = “comparison”, stacked = F, do.stat = TRUE)
gg1 + gg2
~~~
  (B) **Compare outgoing (or incoming) signaling patterns associated with each cell population** CRITICAL The above analysis summarizes the information from the outgoing and incoming signaling together. CellChat can also compare the outgoing (or incoming) signaling pattern between two datasets, allowing to identify signaling pathways/ligand-receptors that exhibit different signaling patterns. CellChat can combine all the identified signaling pathways from different datasets and thus compare them side by side, including outgoing signaling, incoming signaling and overall signaling by aggregating outgoing and incoming signaling together. The ‘`rankNet`’ function also shows the comparison of overall signaling, but it does not show the signaling strength in specific cell populations. CellChat uses heatmap plot to show the contribution of signals (signaling pathways or ligand-receptor pairs) to cell groups in terms of outgoing or incoming signaling. In this heatmap, colobar represents the relative signaling strength of a signaling pathway across cell groups (Note that values are row-scaled). The top colored bar plot shows the total signaling strength of a cell group by summarizing all signaling pathways displayed in the heatmap. The right grey bar plot shows the total signaling strength of a signaling pathway by summarizing all cell groups displayed in the heatmap.

~~~
library(ComplexHeatmap)
i = 1
# combining all the identified signaling pathways from different datasets
pathway.union <- union(object.list[[i]]@netP$pathways, object.list[[i+1]]@netP $pathways)
ht1 = netAnalysis_signalingRole_heatmap(object.list[[i]], pattern = “outgoing “, signaling = pathway.union, title = names(object.list)[i], width = 5, height = 6)
ht2 = netAnalysis_signalingRole_heatmap(object.list[[i+1]], pattern = “outgoin g”, signaling = pathway.union, title = names(object.list)[i+1], width = 5, height = 6)
draw(ht1 + ht2, ht_gap = unit(0.5, “cm”))
~~~ CRITICAL Important parameters of the netAnalysis_signalingRole_heatmap function are as follows:
    - signaling: a character vector giving the name of signaling networks;
    - pattern: “outgoing”, “incoming” or “all”. When pattern = “all”, it aggregates the outgoing and incoming signaling strength together;
    - slot.name: the slot name of object that is used to compute centrality measures of signaling networks;
    - color.use: the character vector defining the color of each cell group;

#### Identify up-regulated and down-regulated signaling ligand-receptor pairs ●Timing ~8 s

22. Identify dysfunctional signaling by comparing the communication probabilities. CellChat can compare the communication probabilities mediated by L-R pairs from certain cell groups to other cell groups. This can be done by setting ‘comparison’ in the function ‘`netVisual_bubble`’. Moreover, CellChat can identify the up-regulated (increased) and down-regulated (decreased) signaling ligand-receptor pairs in one dataset compared to the other dataset. This can be done by specifying ‘max.dataset’ and ‘min.dataset’ in the function ‘`netVisual_bubble`’. The increased signaling means these signaling have higher communication probability (strength) in one dataset compared to the other dataset. The ligand-receptor pairs shown in the bubble plot can be accessed via ‘gg1$data’.

~~~
# Compare the communication probabilities from certain cell groups to other cell groups
netVisual_bubble(cellchat, sources.use = 4, targets.use = c(5:11), comparison = c(1, 2), angle.x = 45)
# Identify the up-regulated (increased) and down-regulated (decreased) signaling ligand-receptor pairs in one dataset compared to the other dataset.
gg1 <- netVisual_bubble(cellchat, sources.use = 4, targets.use = c(5:11), com parison = c(1, 2), max.dataset = 2, title.name = “Increased signaling in LS”, angle.x = 45, remove.isolate = T)
gg2 <- netVisual_bubble(cellchat, sources.use = 4, targets.use = c(5:11), com parison = c(1, 2), max.dataset = 1, title.name = “Decreased signaling in LS”, angle.x = 45, remove.isolate = T)
gg1 + gg2
~~~ CRITICAL Important parameters of the netVisual_bubble function for the comparison analysis are as follows:
  - comparison: a numerical vector giving the datasets for comparison in the merged object; e.g., comparison = c(1,2);
  - group: a numerical vector giving the group information of different datasets; e.g., group = c(1,2,2);
  - max.dataset: a scale, keep the communications with highest probability in max.dataset (i.e., certain condition);
  - min.dataset: a scale, keep the communications with lowest probability in min.dataset;
  - color.text.use whether color the xtick labels according to the dataset origin when doing comparison analysis;
  - color.text the colors for xtick labels according to the dataset origin when doing comparison analysis.
23. Identify dysfunctional signaling by using differential expression analysis. CellChat performs differential expression analysis between two biological conditions (i.e., NL and LS) for each cell group, and then obtain the up-regulated and down-regulated signaling based on the fold change of ligands in the sender cells and receptors in the receiver cells. Such analysis can be done as follows:

~~~
# define a positive dataset, i.e., the dataset with positive fold change against the other dataset
pos.dataset = “LS”
# define a char name used for storing the results of differential expression analysis
features.name = pos.dataset
# perform differential expression analysis
cellchat <- identifyOverExpressedGenes(cellchat, group.dataset = “datasets”, p os.dataset = pos.dataset, features.name = features.name, only.pos = FALSE, thr esh.pc = 0.1, thresh.fc = 0.1, thresh.p = 1)
# map the results of differential expression analysis onto the inferred cell-cell communications to easily manage/subset the ligand-receptor pairs of interest
net <- netMappingDEG(cellchat, features.name = features.name)
# extract the ligand-receptor pairs with upregulated ligands in LS
net.up <- subsetCommunication(cellchat, net = net, datasets = “LS”,ligand.logFC = 0.2, receptor.logFC = NULL)
# extract the ligand-receptor pairs with upregulated ligands and upregulated r ecetptors in NL, i.e.,downregulated in LS
net.down <- subsetCommunication(cellchat, net = net, datasets = “NL”,ligand.logFC = −0.1, receptor.logFC = −0.1)
# do further deconvolution to obtain the individual signaling genes
gene.up <- extractGeneSubsetFromPair(net.up, cellchat)
gene.down <- extractGeneSubsetFromPair(net.down, cellchat)
# Users can also find all the significant outgoing/incoming/both signaling acc ording to the customized DEG features and cell groups of interest
df <- findEnrichedSignaling(object.list[[2]], features = c(“CCL19”, “CXCL12 “), idents = c(“Inflam. FIB”, “COL11A1+ FIB”), pattern =“outgoing”)
~~~
24. Visualize the identified up-regulated and down-regulated signaling ligand-receptor pairs using bubble plot (option A), chord diagram (option B) or wordcloud (option C).
  (A) **Bubble plot**

~~~
pairLR.use.up = net.up[, “interaction_name”, drop = F]
gg1 <- netVisual_bubble(cellchat, pairLR.use = pairLR.use.up, sources.use = 4, targets.use = c(5:11), comparison = c(1, 2), angle.x = 90, remove.isolate = T,title.name = paste0(“Up-regulated signaling in “, names(object.list)[2]))
pairLR.use.down = net.down[, “interaction_name”, drop = F]
gg2 <- netVisual_bubble(cellchat, pairLR.use = pairLR.use.down, sources.use = 4, targets.use = c(5:11), comparison = c(1, 2), angle.x = 90, remove.isolate = T,title.name = paste0(“Down-regulated signaling in “, names(object.list)[2]))
gg1 + gg2
~~~
  (B) **Chord diagram**

~~~
par(mfrow = c(1,2), xpd=TRUE)
netVisual_chord_gene(object.list[[2]], sources.use = 4, targets.use = c(5:11), slot.name = ‘net’, net = net.up, lab.cex = 0.8, small.gap = 3.5, title.name = paste0(“Up-regulated signaling in “, names(object.list)[2]))
netVisual_chord_gene(object.list[[1]], sources.use = 4, targets.use = c(5:11), slot.name = ‘net’, net = net.down, lab.cex = 0.8, small.gap = 3.5, title.name = paste0(“Down-regulated signaling in “, names(object.list)[2]))
~~~
  (C) **Wordcloud plot**

~~~
# visualize the enriched ligands in the first condition
computeEnrichmentScore(net.down, species = ‘human’)
# visualize the enriched ligands in the second condition
computeEnrichmentScore(net.up, species = ‘human’)
~~~

#### Visually compare inferred cell-cell communication networks ●Timing ~5 s

25. Visualize inferred cell-cell communication networks. Similar to CellChat analysis of individual dataset (**Part 1**), CellChat can visualize cell-cell communication network using hierarchy plot, circle plot, chord diagram, or heatmap. Here we briefly show two examples, including circle plot (option A) and heatmap plot (option B). More details on the visualization can be found in Step 9.
  (A) **Circle plot**

~~~
pathways.show <- c(“CXCL”)
weight.max <- getMaxWeight(object.list, slot.name = c(“netP”), attribute = pathways.show) # control the edge weights across different datasets
par(mfrow = c(1,2), xpd=TRUE)
for (i in 1:length(object.list)) {
netVisual_aggregate(object.list[[i]], signaling = pathways.show, layout = “c ircle”, edge.weight.max = weight.max[1], edge.width.max = 10, signaling.name = paste(pathways.show, names(object.list)[i]))
}
~~~
  (B) **Heatmap plot**

~~~
pathways.show <- c(“CXCL”)
par(mfrow = c(1,2), xpd=TRUE)
ht <- list()
for (i in 1:length(object.list)) {
 ht[[i]] <- netVisual_heatmap(object.list[[i]], signaling = pathways.show, color.heatmap = “Reds”,title.name = paste(pathways.show, “signaling “,names(object.list)[i]))
}
ComplexHeatmap::draw(ht[[1]] + ht[[2]], ht_gap = unit(0.5, “cm”))
~~~
26. Visualize gene expression distribution. CellChat can plot the gene expression distribution of signaling genes related to L-R pairs or signaling pathway using a Seurat wrapper function ‘`plotGeneExpression`’.

~~~
cellchat@meta$datasets = factor(cellchat@meta$datasets, levels = c(“NL”, “LS “)) # set factor level
plotGeneExpression(cellchat, signaling = “CXCL”, split.by = “datasets”, color s.ggplot = T)
~~~ ◼ PAUSE POINT Users can now export the merged CellChat object and the list of the two separate objects for later use:

~~~
save(object.list, file = “cellchat_object.list_humanSkin_NL_LS.RData”)
save(cellchat, file = “cellchat_merged_humanSkin_NL_LS.RData”)
~~~

#### Part 3. Comparison analysis of multiple datasets with differing cell type compositions ● Timing ~5 s

▴ CRITICAL In this section, we demonstrate how to apply CellChat to the comparative analysis of multiple conditions with differing cell type compositions. For the datasets with differing cell type (group) compositions, CellChat can lift up the cell groups to the same cell labels across all datasets using the function ‘`liftCellChat`’, and then perform comparative analysis as the joint analysis of datasets with the same cell type compositions. The equivalent online version of this described procedure (Steps 27-29) along with the graphical plots are available in the tutorial directory of the CellChat github repository (https://htmlpreview.github.io/?https://github.com/jinworks/CellChat/blob/master/tutorial/Comparison_analysis_of_multiple_datasets_with_different_cellular_compositions.html). Here we take an example of comparative analysis of two embryonic mouse skin scRNA-seq datasets from days E13.5 and E14.5. There are 11 shared skin cell populations at E13.5 and E14.5 and additional two populations (i.e., dermal DC and pericytes) specific to E14.5. Therefore, we lift up the cell groups from E13.5 to the same cell labels as E14.5.

27. Load CellChat object of each dataset. Users need to run CellChat on each dataset separately and then merge different CellChat objects together. Here we also do ‘updateCellChat’ because these two objects are obtained using the earlier version (< 1.6.0) of CellChat.

~~~
cellchat.E13 <- readRDS(“./tutorial/cellchat_embryonic_E13.rds”)
cellchat.E13 <- updateCellChat(cellchat.E13)
cellchat.E14 <- readRDS(“./tutorial/cellchat_embryonic_E14.rds”)
cellchat.E14 <- updateCellChat(cellchat.E14)
~~~
28. Lift up CellChat objects and merge them together. Since there are two additional populations (i.e., dermal DC and pericytes) specific to E14.5 compared to E13.5, we lift up ‘cellchat.E13’ by lifting up the cell groups to the same cell labels as E14.5. ‘liftCellChat’ only updates the slot related to cell-cell communication network, including slots object@net, object@netP and object@idents.

~~~
group.new = levels(cellchat.E14@idents) # Define the cell labels to lift up
cellchat.E13 <- liftCellChat(cellchat.E13, group.new)
object.list <- list(E13 = cellchat.E13, E14 = cellchat.E14)
cellchat <- mergeCellChat(object.list, add.names = names(object.list), cell.prefix = TRUE)
~~~
29. Comparative visualization and analysis of cell-cell communication. Once we lift up the CellChat object and merge different objects together, we can perform CellChat analysis the same way as the comparative analysis of multiple datasets with the same cell type compositions (see Steps 17-26 in **Part 2**). Below is an example of how to compare the inferred cell-cell communication networks using circle plot.

~~~
# Circle plot
pathways.show <- c(“WNT”)
weight.max <- getMaxWeight(object.list, slot.name = c(“netP”), attribute = pat hways.show) # control the edge weights across different datasets
par(mfrow = c(1,2), xpd=TRUE)
for (i in 1:length(object.list)) {
 netVisual_aggregate(object.list[[i]], signaling = pathways.show, layout = “c ircle”, edge.weight.max = weight.max[1], edge.width.max = 10, signaling.name = paste(pathways.show, names(object.list)[i]))
}
~~~ ◼PAUSE POINT Users can now export the merged CellChat object and the list of the two separate objects for later use:

~~~
save(object.list, file = “cellchat_object.list_embryonic_E13_E14. RData”)
save(cellchat, file = “cellchat_merged_embryonic_E13_E14.RData”)
~~~

#### Part 4. Inferring spatially-proximal cell-cell communication from spatially resolved transcriptomic data

▴ CRITICAL In this section, we demonstrate CellChat’s ability to infer, analyze and visualize spatially-proximal cell-cell communication network from a single spatially resolved transcriptomic dataset. CellChat requires gene expression and spatial location data of spots/cells as the user input and models the probability of cell-cell communication by integrating gene expression with spatial distance as well as prior knowledge of the interactions between signaling ligands, receptors and their cofactors. The detailed descriptions of the input files for inferring spatially-proximal cell-cell communication can be found in the section **Required input data**.

#### Data input & preprocessing from spatially resolved transcriptomic data ●Timing ~15 s

30. Prepare input data. Example dataset is a mouse brain 10X Visium dataset (https://www.10xgenomics.com/resources/datasets/mouse-brain-serial-section-1-sagittal-anterior-1-standard-1-0-0). Biological annotations of spots (i.e., cell group information) are predicted using Seurat R package^46^ (Fig. 4b). The equivalent online version of this described procedure (Steps 30-35) along with the graphical plots are available in the tutorial directory of the CellChat github repository (https://htmlpreview.github.io/?https://github.com/jinworks/CellChat/blob/master/tutorial/CellChat_analysis_of_spatial_imaging_data.html). We then prepare input data for CellChat analysis:

~~~
library(CellChat)
library(patchwork)
library(Seurat)
load(“./tutorial/visium_mouse_cortex_annotated.RData”)
# Gene expression data
data.input = GetAssayData(visium.brain, slot = “data”, assay = “SCT”) # normalized data matrix
# User assigned cell labels
meta = data.frame(labels = Idents(visium.brain), row.names = name s(Idents(visium.brain)))
# Spatial locations of spots from full (NOT high/low) resolution images
spatial.locs = GetTissueCoordinates(visium.brain, scale = NULL, cols = c(“imagerow”, “imagecol”))
# Scale factors and spot diameters of the full resolution images
scale.factors = jsonlite::fromJSON(txt = file.path(“./tutorial/sp atial_imaging_data_visium-brain”, ‘scalefactors_json.json’))
scale.factors = list(spot.diameter = 65, spot = scale.factors$spot_diameter_fullres)
~~~ CRITICAL With the digital gene expression matrix, cell label information and spatial imaging information, we can follow the same steps as the conventional CellChat analysis of non-spatial scRNA-seq data (see **Part 1**). Below we briefly go through the key steps for inferring spatially-proximal cell-cell communication.
31. Create a CellChat object. We initialize a CellChat object by typing the following:

~~~
cellchat <- createCellChat(object = data.input, meta = meta, group.by = “labels”, datatype = “spatial”, coordinates = spatial.locs, scale.factors = scale.factors)
~~~ CRITICAL Spatially-related parameters of the `createCellChat` function are as follows:
  - datatype: By default datatype = “RNA”; when running CellChat on spatial imaging data, set datatype = “spatial” and input ‘scale.factors’;
  - coordinates: a data matrix in which each row gives the spatial locations/coordinates of each cell/spot;
  - scale.factors: a list containing the scale factors and spot diameter for the full resolution images. User must input this list when datatype = “spatial”. scale.factors must contain an element named ‘spot.diameter’, which is the theoretical spot size; e.g., 10x Visium (spot.size = 65 *μm*), and another element named ‘spot’, which is the number of pixels that span the diameter of a theoretical spot size in the original, full-resolution image. It should be noted that the parameter ‘spot.diameter’ is dependent on spatial imaging technologies and parameter ‘spot’ is dependent on specific datasets.
32. Set ligand-receptor interaction database.

~~~
CellChatDB <- CellChatDB.mouse
# use a subset of CellChatDB for cell-cell communication analysis
CellChatDB.use <- subsetDB(CellChatDB, search = “Secreted Signaling”) # use Secreted Signaling
# CellChatDB.use <- CellChatDB # simply use the default CellChatDB to use all CellChatDB for cell-cell communication analysis
# set the used database in the object
cellchat@DB <- CellChatDB.use
~~~
33. Identify over-expressed ligands or receptors.

~~~
cellchat <- subsetData(cellchat) # This step is necessary even if using the whole database
future::plan(“multisession”, workers = 4) # do parallel
cellchat <- identifyOverExpressedGenes(cellchat)
cellchat <- identifyOverExpressedInteractions(cellchat)
~~~

#### Inference of cell-cell communication network from spatially resolved transcriptomic data

●Timing ~15 min

34. Infer cell-cell communication network at both ligand-receptor pair and signaling pathway levels.

~~~
cellchat <- computeCommunProb(cellchat, type = “truncatedMean”, trim = 0.1,
                  distance.use = TRUE, interaction.range = 250, scale.distance = 0.01)
# Filter the cell-cell communication
cellchat <- filterCommunication(cellchat, min.cells = 10)
# Infer the cell-cell communication at a signaling pathway level
cellchat <- computeCommunProbPathway(cellchat)
# Calculate the aggregated cell-cell communication network
cellchat <- aggregateNet(cellchat)
~~~ CRITICAL Spatially-related parameters of the `computeCommunProb` function are as follows:
  - distance.use: whether to use distance constraints to compute communication probability. By default, distance.use = TRUE. When setting distance.use = FALSE, it only filters out interactions between spatially distant regions, but not add distance constraints;
  - interaction.range: the maximum interaction/diffusion range of ligands (Unit: *μm*). This hard threshold is used to filter out the connections between spatially distant regions;
  - scale.distance: a scale or normalization factor for the spatial distances. This value can be 1, 0.1, 0.01, 0.001. We choose this value such that the minimum value of the scaled distances is in [1,2]. When comparing communication across different CellChat objects, the same scale factor should be used. For a single CellChat analysis, different scale factors do not affect the ranking of the signaling based on their interaction strength. USERS may need to adjust the parameter ‘scale.distance’ when working on data from other spatial imaging technologies;
  - k.min: the minimum number of interacting cell pairs required for defining adjacent cell groups.

◼PAUSE POINT Users can now export the merged CellChat object for later use:

~~~
saveRDS(cellchat, file = “cellchat_visium_mouse_cortex.Rds”)
~~~

### Visualization and analysis of cell-cell communication network in a spatial context ●Timing ~1 s

35. Upon inferring the intercellular communication network from spatial imaging data, CellChat’s various functionality can be used for further data exploration, analysis, and visualization. Please check more in **Part 1**. Here we only showcase the ‘circle plot’ and the newly added ‘spatial plot’. We first show the number of interactions or the total interaction strength (weights) between any two cell groups using circle plot (option A), and then visualize the cell-cell communication network using circle plot and spatial plot (option B).
  (A) **Visualize the number of interactions or interaction strength between any two cell groups**

~~~
groupSize <- as.numeric(table(cellchat@idents))
par(mfrow = c(1,2), xpd=TRUE)
netVisual_circle(cellchat@net$count, vertex.weight = rowSums(cell chat@net$count), weight.scale = T, label.edge= F, title.name = “Number of interactions”)
netVisual_circle(cellchat@net$weight, vertex.weight = rowSums(cel lchat@net$weight), weight.scale = T, label.edge= F, title.name = “Interaction weights/strength”)
~~~
  (B) **Visualize the inferred cell-cell communication network of a specific signaling pathway**

~~~
pathways.show <- c(“CXCL”)
 # Circle plot
 par(mfrow=c(1,1))
 netVisual_aggregate(cellchat, signaling = pathways.show, layout = “circle”)
 # Spatial plot
 par(mfrow=c(1,1))
 netVisual_aggregate(cellchat, signaling = pathways.show, layout = “spatial”, edge.width.max = 2, vertex.size.max = 1, alpha.image = 0.2, vertex.label.cex = 3.5).
~~~

### Timing

- Steps 1-4, data input & preprocessing: ∼12 s;
- Steps 5-8, inference of cell-cell communication network: ∼37 s;
- Steps 9-12, visualization of cell-cell communication network: ∼10 s;
- Steps 13-15, systems analysis of cell-cell communication network: ∼2.7 mins.
- Step 16, load CellChat object of each dataset and merge them together: ∼3 s;
- Steps 17-19, predict general principles of cell-cell communication: ∼2 s;
- Steps 20-21, identify altered signaling with distinct network architecture and interaction strength: ∼17 s;
- Steps 22-26, identify up-regulated and down-regulated signaling ligand-receptor pairs:∼8 s;
- Steps 27-29, comparison analysis of multiple datasets with differing cell type compositions: ∼5 s.
- Steps 30-33, data input & preprocessing from spatially resolved transcriptomic data:∼15 s;
- Step 34, inference of cell-cell communication network from spatially resolved transcriptomic data: ∼15 min;
- Step 35, visualization and analysis of cell-cell communication network in a spatial context: ∼1 s.

### Troubleshooting

Troubleshooting advice can be found in Table 1.

**Table 1.**
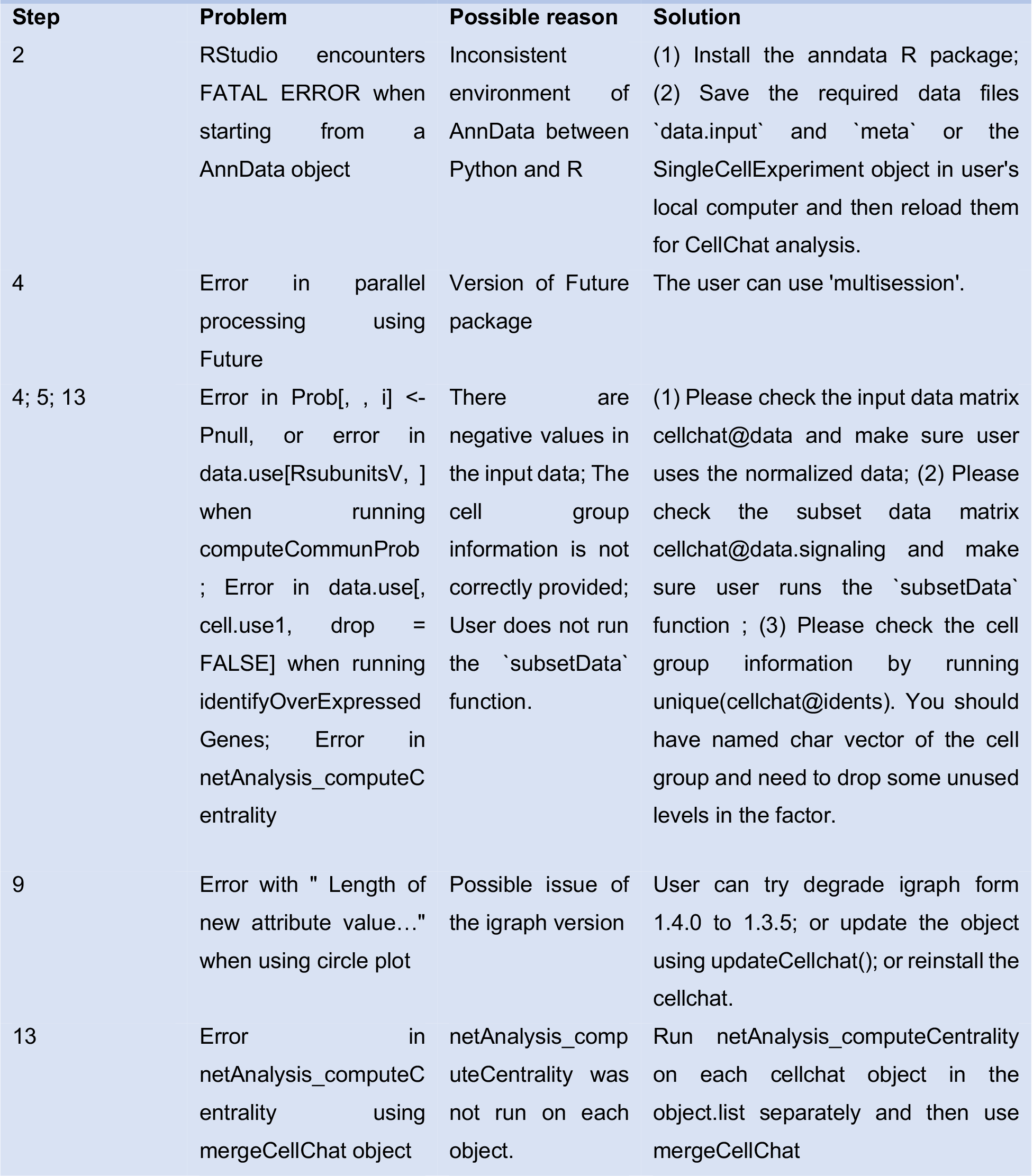
Troubleshooting table.

### Anticipated results

Running CellChat’s inference (Step 5) of the ligand-receptor pair-mediated cell-cell communication produces the communication probability (i.e., interaction strength) array and the corresponding p-value array, which can be accessed by object@net$prob and object@net$pval, respectively. Both arrays are three dimensional, where the first, second and third dimensions represent sources, targets and ligand-receptor pairs, respectively (Fig. 1). For example, given an inferred cell-cell communication probability array *P*(*x,y, z*), *P*(*x*_*i*_,*y*_*j*_, *z*_*k*_) is the communication probability from cell group *x*_*i*_ to cell group *y*_*j*_ for a ligand-receptor pair *z*_*k*_. CellChat also infers signaling pathway-mediated cell-cell communication (Step 7), where the communication probability array can be accessed by object@netP$prob. CellChat provides several ways to visualize the inferred cell-cell communication network, including circle plot, hierarchical plot, chord diagram, heatmap, spatial plot and bubble plot. Of note, users can visualize inferred communication networks of an individual L-R pair, a signaling pathway as well as multiple ligand-receptor pairs or signaling pathways. To facilitate the interpretation of the inferred intercellular communication networks within one condition and across different conditions (Fig. 1), CellChat can (i) identify signaling roles of cell groups as well as the major contributing signaling within a given signaling network; (ii) predict key incoming and outgoing signals for specific cell types as well as global communication patterns on how multiple cell types and signaling pathways coordinate together; (iii) group signaling pathways from both functional and topological perspectives; (iv) identify major signaling changes and altered cell populations across different biological conditions using various quantitative metrics and differential expression analysis; and (v) perform comparison analysis across different conditions with differing cell type compositions.

The inferred cell-cell communications depend on the method for computing average expression per cell group. The “triMean” method produces fewer but stronger interactions, while the “truncatedMean” method with a smaller value of the “trim” parameter (e.g., ‘trim = 0.1’) produces more interactions, leading to the detection of weak signaling. To demonstrate this point, we compare the number of inferred interactions and the enriched signaling pathways when using “triMean”, “truncatedMean” with ‘trim = 0.1’, and “truncatedMean” with ‘trim = 0.05’, respectively (Figs. 3a and 3b). Therefore, if known signaling is not observed, users can use ‘truncatedMean’ with lower values of ‘trim’ to change the method for calculating the average gene expression per cell group.

When spatial locations of cells are available, CellChat v2 further infers spatially proximal cell-cell communication between cell groups (see the Development of the protocol section and Fig. 4a on this new method in detail). In this protocol (Steps 30-35), we applied our new CellChat v2 to a mouse brain 10X Visium dataset, and compared the inferred results with those obtained from the original CellChat without spatial constraints (Figs. 4b-d). By comparing the number of interactions between pairs of cell groups (Fig. 4c), we observe that CellChat v2 is able to filter out the interactions between spatially distant groups of cells. In addition, CellChat v2 offers new visualization outputs in the spatial context (Fig. 4d), including spatial plot for visualizing the inferred interactions and feature plot for examining the expression distribution of ligands and receptors on the tissue image. We also applied CellChat v2 to another 10X Visium spatial transcriptomic data of human adult intestine, which is less structured across 25 cell subpopulations compared to brain tissue (Supplementary Fig. S1). Similarly, CellChat v2 is able to filter out interactions between spatially distant groups of cells.

Finally, CellChat v2 allows users to visualize and explore the data and the inferred signaling interactively by defining various analysis parameters (Fig. 5). Briefly, it can visualize cell groups and signaling expression on the tissue, examine the inferred signaling between different cell groups and further visualize the individual signaling pathway. A rich user-guided sliders in each panel are provided for flexible exploration.

**Figure 5:**
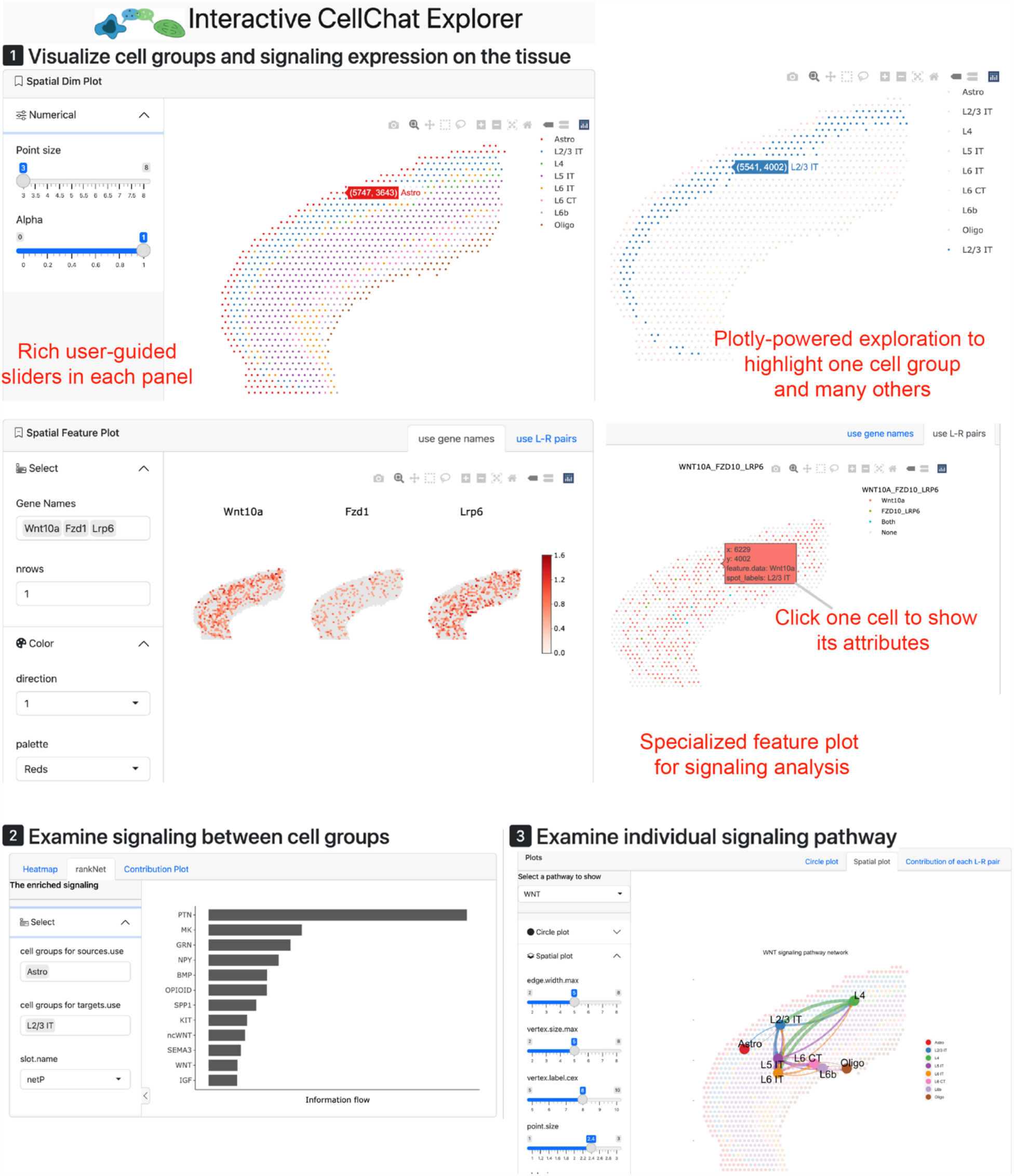
Overview of the Interactive CellChat Explorer created by runCellChatApp function in the R package. To facilitate the exploration of spatially proximal cell-cell communication, CellChat allows the end-user to visualize and explore the data and the inferred signaling interactively. It can visualize cell groups and signaling expression on the tissue, examine the inferred signaling between different cell groups and further visualize the individual signaling pathway. A rich user-guided sliders in each panel are provided for flexible exploration, highlight and zoom-out of the related information of interest.

### Reporting Summary

Further information on research design is available in the Nature Research Reporting Summary linked to this article.

## Data availability

The example datasets analyzed in this protocol are all publicly available. The data used in Steps 1-15 are available at the Gene Expression Omnibus (GEO) under accession GSE147424^47^. The data used in Steps 16-26 are available at the GEO under accession GSM3453535, GSM3453536, GSM3453537, and GSM3453538^48^. The data used in Steps 30-35 are available at https://www.10xgenomics.com/resources/datasets/mouse-brain-serial-section-1-sagittal-anterior-1-standard-1-0-0). The human intestine data were obtained from https://simmonslab.shinyapps.io/FetalAtlasDataPortal/. In addition, all the preprocessed datasets and CellChat objects required to reproduce this protocol are publicly available at https://figshare.com/projects/Example_data_for_cell-cell_communication_analysis_using_CellChat/157272.

## Code availability

CellChat is publicly available as an R package. Source codes of the R package and this protocol have been deposited at the GitHub repository (https://github.com/jinworks/CellChat).

## Author contributions statements

SJ and QN designed the project, SJ implemented the codes and developed the protocol, SJ and QN drafted the manuscript, SJ, QN and MVP edited and approved the manuscript.

## Acknowledgments

The work was partly supported by a National Science Foundation grants DMS1763272, MCB2028424, and CBET2134916, a grant from the Simons Foundation (594598 to Q.N.), National Institutes of Health grants R01AR079150 and R01DE030565, and a Chan Zuckerberg Initiative grant (AN-0000000062). The authors thank Ye Su in the School of Mathematics and Statistics at Wuhan University for the help on implementing the R Shiny app.

## Competing interests

The authors declare no competing interests.

Key references using this protocol

## Figures

### Box 1

Updating the ligand-receptor interaction database CellChatDB

Users can use the function ‘`updateCellChatDB`’ to update the ligand-receptor interaction database CellChatDB by integrating new ligand-receptor pairs from other resources or utilizing a custom ligand-receptor interaction database. The minimum input of this function is a data frame with two columns named as ‘ligand’ and ‘receptor’, and other inputs include classified pathway, complex, cofactor and gene information. Users can check the documentation of this function for more details of the required input data or the online tutorial (https://htmlpreview.github.io/?https://github.com/jinworks/CellChat/blob/master/tutorial/Update-CellChatDB.html). Here we describe how to integrate other resources by taking another database CellTalkDB as an example. After downloading the CellTalkDB from https://github.com/ZJUFanLab/CellTalkDB, type the following:

~~~
# Load the customized ligand-receptor pairs: a data frame with two columns
named as ‘ligand’ and ‘receptor’
db.user <- readRDS(“./CellTalkDB-master/database/human_lr_pair.rds”)
# Load the gene information: a data frame with one column named as ‘Symbol’.
gene_info <- readRDS(“./CellTalkDB-master/data/human_gene_info.rds”)
# Modify the colnames if needed
colnames(db.user) <- plyr::mapvalues(colnames(db.user), from = c(“ligand_gene_symbol”,”receptor_gene_symbol”,”lr_pair”), to = c(“ligand”,”receptor”,”interaction_name”), warn_missing = TRUE)
# Create a new database by typing one of the following commands
db.new <- updateCellChatDB(db = db.user, gene_info = gene_info)
db.new <- updateCellChatDB(db = db.user, gene_info = NULL, species_target = “human”)
# Alternatively, users can integrate the customized L-R pairs into the built-in CellChatDB
db.new <- updateCellChatDB(db = db.user, merged = TRUE, species_target = “human”)
# Users can now use this new database in CellChat analysis (Step 3 of this protocol)
cellchat@DB <- db.new
# Users can save the new database for future use
save(db.new, file = “CellChatDB.human_user.rda”)
~~~

## Notes

https://github.com/jinworks/CellChat

### Competing Interest Statement

The authors have declared no competing interest.

## References

1. Shao, X., Lu, X., Liao, J., Chen, H. & Fan, X. New avenues for systematically inferring cell-cell communication: through single-cell transcriptomics data. Protein Cell 11, 866–880 (2020).

2. Armingol, E., Officer, A., Harismendy, O. & Lewis, N.E. Deciphering cell-cell interactions and communication from gene expression. Nat Rev Genet 22, 71–88 (2021).

3. Almet, A.A., Cang, Z., Jin, S. & Nie, Q. The landscape of cell–cell communication through single-cell transcriptomics. Current Opinion in Systems Biology 26, 12–23 (2021).

4. Longo, S.K., Guo, M.G., Ji, A.L. & Khavari, P.A. Integrating single-cell and spatial transcriptomics to elucidate intercellular tissue dynamics. Nat Rev Genet 22, 627–644 (2021).

5. Jin, S. & Ramos, R. Computational exploration of cellular communication in skin from emerging single-cell and spatial transcriptomic data. Biochem Soc Trans 50, 297–308 (2022).

6. Wang, X., Almet, A.A. & Nie, Q. The promising application of cell-cell interaction analysis in cancer From single-cell and spatial transcriptomics. Semin Cancer Biol (2023).

7. Jin, S. et al. Inference and analysis of cell-cell communication using CellChat. Nat Commun 12, 1088 (2021).

8. Vu, R. et al. Wound healing in aged skin exhibits systems-level alterations in cellular composition and cell-cell communication. Cell Rep 40, 111155 (2022).

9. Hao, M., Zou, X. & Jin, S. Identification of Intercellular Signaling Changes Across Conditions and Their Influence on Intracellular Signaling Response From Multiple Single-Cell Datasets. Front Genet 12, 751158 (2021).

10. Kanemaru, K. et al. Spatially resolved multiomics of human cardiac niches. Nature 619, 801–810 (2023).

11. Zhao, W., Johnston, K.G., Ren, H., Xu, X. & Nie, Q. Inferring neuron-neuron communications from single-cell transcriptomics through NeuronChat. Nat Commun 14, 1128 (2023).

12. Yang, A.C. et al. Dysregulation of brain and choroid plexus cell types in severe COVID-19. Nature 595, 565–571 (2021).

13. Wendisch, D. et al. SARS-CoV-2 infection triggers profibrotic macrophage responses and lung fibrosis. Cell 184, 6243–6261 e6227 (2021).

14. Winkler, E.A. et al. A single-cell atlas of the normal and malformed human brain vasculature. Science 375, eabi7377 (2022).

15. Lake, B.B. et al. An atlas of healthy and injured cell states and niches in the human kidney. Nature 619, 585–594 (2023).

16. Cheng, Y.T. et al. Inhibitory input directs astrocyte morphogenesis through glial GABA(B)R. Nature 617, 369–376 (2023).

17. Ortiz-Munoz, G. et al. In situ tumour arrays reveal early environmental control of cancer immunity. Nature 618, 827–833 (2023).

18. Li, J. et al. Remodeling of the immune and stromal cell compartment by PD-1 blockade in mismatch repair-deficient colorectal cancer. Cancer Cell 41, 1152–1169 e1157 (2023).

19. Lim, X. & Nusse, R. Wnt signaling in skin development, homeostasis, and disease. Cold Spring Harb Perspect Biol 5 (2013).

20. Dimitrov, D. et al. Comparison of methods and resources for cell-cell communication inference from single-cell RNA-Seq data. Nat Commun 13, 3224 (2022).

21. Liu, Z., Sun, D. & Wang, C. Evaluation of cell-cell interaction methods by integrating single-cell RNA sequencing data with spatial information. Genome Biol 23, 218 (2022).

22. Efremova, M., Vento-Tormo, M., Teichmann, S.A. & Vento-Tormo, R. CellPhoneDB: inferring cell-cell communication from combined expression of multi-subunit ligand-receptor complexes. Nat Protoc 15, 1484–1506 (2020).

23. Garcia-Alonso, L. et al. Mapping the temporal and spatial dynamics of the human endometrium in vivo and in vitro. Nat Genet 53, 1698–1711 (2021).

24. Hou, R., Denisenko, E., Ong, H.T., Ramilowski, J.A. & Forrest, A.R.R. Predicting cell-to-cell communication networks using NATMI. Nat Commun 11, 5011 (2020).

25. Noel, F. et al. Dissection of intercellular communication using the transcriptome-based framework ICELLNET. Nat Commun 12, 1089 (2021).

26. Raredon, M.S.B. et al. Computation and visualization of cell-cell signaling topologies in single-cell systems data using Connectome. Sci Rep 12, 4187 (2022).

27. Armingol, E. et al. Context-aware deconvolution of cell-cell communication with Tensor-cell2cell. Nat Commun 13, 3665 (2022).

28. Luo, J., Deng, M., Zhang, X. & Sun, X. ESICCC as a systematic computational framework for evaluation, selection, and integration of cell-cell communication inference methods. LID - gr.278001.123 [pii] LID - 10.1101/gr.278001.123 [doi]. (2023).

29. Vento-Tormo, R. et al. Single-cell reconstruction of the early maternal-fetal interface in humans. Nature 563, 347–353 (2018).

30. Garcia-Alonso, L. et al. Single-cell roadmap of human gonadal development. Nature 607, 540–547 (2022).

31. Browaeys, R. et al. MultiNicheNet: a flexible framework for differential cell-cell communication analysis from multi-sample multi-condition single-cell transcriptomics data. bioRxiv (2023).

32. Cang, Z. & Nie, Q. Inferring spatial and signaling relationships between cells from single cell transcriptomic data. Nat Commun 11, 2084 (2020).

33. Shao, X. et al. Knowledge-graph-based cell-cell communication inference for spatially resolved transcriptomic data with SpaTalk. Nat Commun 13, 4429 (2022).

34. Cang, Z. et al. Screening cell-cell communication in spatial transcriptomics via collective optimal transport. Nat Methods 20, 218–228 (2023).

35. Shao, X. et al. CellTalkDB: a manually curated database of ligand-receptor interactions in humans and mice. Brief Bioinform 22, bbaa269 (2021).

36. Perkel, J.M. Single-cell proteomics takes centre stage. Nature 597, 580–582 (2021).

37. Jin, S., Zhang, L. & Nie, Q. scAI: an unsupervised approach for the integrative analysis of parallel single-cell transcriptomic and epigenomic profiles. Genome Biol 21, 25 (2020).

38. Zhang, L., Zhang, J. & Nie, Q. DIRECT-NET: An efficient method to discover cisregulatory elements and construct regulatory networks from single-cell multiomics data. Sci Adv 8, eabl7393 (2022).

39. Wang, S., Karikomi, M., MacLean, A.L. & Nie, Q. Cell lineage and communication network inference via optimization for single-cell transcriptomics. Nucleic Acids Res 47, e66 (2019).

40. Browaeys, R., Saelens, W. & Saeys, Y. NicheNet: modeling intercellular communication by linking ligands to target genes. Nat Methods 17, 159–162 (2020).

41. Hu, Y., Peng, T., Gao, L. & Tan, K. CytoTalk: De novo construction of signal transduction networks using single-cell transcriptomic data. Sci Adv 7 (2021).

42. Zhang, Y. et al. CellCall: integrating paired ligand-receptor and transcription factor activities for cell-cell communication. Nucleic Acids Res 49, 8520–8534 (2021).

43. Cheng, J., Zhang, J., Wu, Z. & Sun, X. Inferring microenvironmental regulation of gene expression from single-cell RNA sequencing data using scMLnet with an application to COVID-19. Brief Bioinform 22, 988–1005 (2021).

44. Zhuang, X. Spatially resolved single-cell genomics and transcriptomics by imaging. Nat Methods 18, 18–22 (2021).

45. Landherr, A., Friedl, B. & Heidemann, J. A critical review of centrality measures in social networks. Business & Information Systems Engineering 2, 371–385 (2010).

46. Stuart, T. et al. Comprehensive integration of single-cell data. Cell 177, 1888–1902 e1821 (2019).

47. He, H. et al. Single-cell transcriptome analysis of human skin identifies novel fibroblast subpopulation and enrichment of immune subsets in atopic dermatitis. J Allergy Clin Immunol 145, 1615–1628 (2020).

48. Gupta, K. et al. Single-Cell Analysis Reveals a Hair Follicle Dermal Niche Molecular Differentiation Trajectory that Begins Prior to Morphogenesis. Dev Cell 48, 17–31 e16 (2019).

## Related links

Jin, S. et al. Nat Commun 12, 1088 (2021): 10.1038/s41467-021-21246-9

Vu, R. & Jin, S. et al. Cell Rep 40, 111155 (2022): 10.1016/j.celrep.2022.111155

