## Supplementary Figure S1 for "CellChat for systematic analysis of cell-cell communication from single-cell and spatially resolved transcriptomics"

a

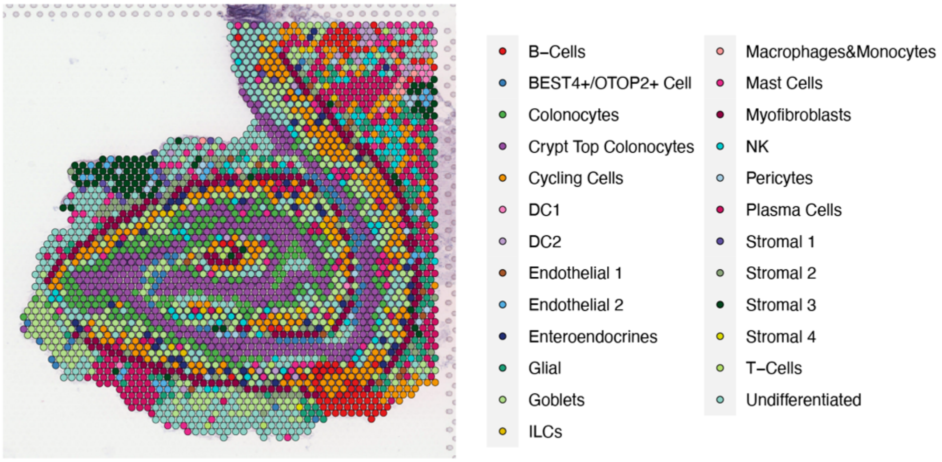


b

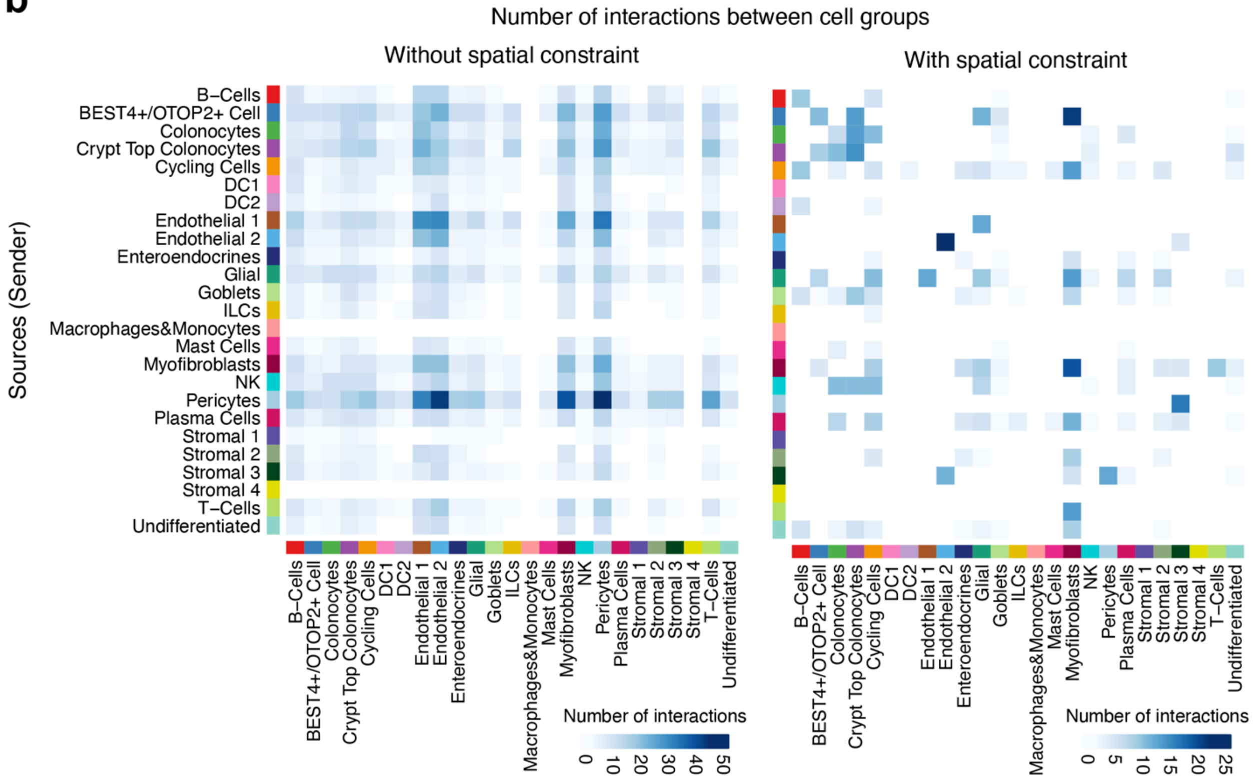


**Supplementary Figure S1: Application of CellChat v2 to human intestine spatially resolved transcriptomics. (a)** Annotations of each spot are used as cell group information for CellChat analysis. **(b)** Comparison of the number of interactions between pairs of cell groups using spatial constraint against the case without spatial constraint. Rows are sources acting as senders, and columns are targets acting as receivers.
